# The Impact of Scene Context on Visual Object Recognition: Comparing Humans, Monkeys, and Computational Models

**DOI:** 10.1101/2024.05.27.596127

**Authors:** Sara Djambazovska, Anaa Zafer, Hamidreza Ramezanpour, Gabriel Kreiman, Kohitij Kar

## Abstract

During natural vision, we rarely see objects in isolation but rather embedded in rich and complex contexts. Understanding how the brain recognizes objects in natural scenes by integrating contextual information remains a key challenge. To elucidate neural mechanisms compatible with human visual processing, we need an animal model that behaves similarly to humans, so that inferred neural mechanisms can provide hypotheses relevant to the human brain. Here we assessed whether rhesus macaques could model human context-driven object recognition by quantifying visual object identification abilities across variations in the amount, quality, and congruency of contextual cues. Behavioral metrics revealed strikingly similar context-dependent patterns between humans and monkeys. However, neural responses in the inferior temporal (IT) cortex of monkeys that were never explicitly trained to discriminate objects in context, as well as current artificial neural network models, could only partially explain this cross-species correspondence. The shared behavioral variance unexplained by context-naive neural data or computational models highlights fundamental knowledge gaps. Our findings demonstrate an intriguing alignment of human and monkey visual object processing that defies full explanation by either brain activity in a key visual region or state-of-the-art models.

## Introduction

The field of visual neuroscience has long been fascinated by the computationally remarkable process of object recognition^1–3^, a cornerstone of primate visual perception. However, understanding an image transcends the ability to identify specific and isolated objects^4–6^. Interpreting an image requires knowledge about object correlations (e.g., bananas tend to co-occur with trees), relative object sizes (e.g., bananas are often smaller than trees), and relative object positions (e.g., bananas tend to be near the top part of a tree). Contextual information can dramatically alter how object information is interpreted ^7,8^. There has been a long-standing interest in the statistics of natural images, and there are foundational behavioral studies of the role of context in vision ^9–14^. The mechanisms behind incorporating contextual cues at the computational and neurophysiological levels remain poorly understood. Multiple prior studies focused on the role of context in relatively “low-level” visual phenomena such as extra-classical receptive fields and surround suppression ^15–19^. However, little is known about how the brain represents prior high-level knowledge and integrates it with incoming inputs to modulate visual cognition.

Over the last decades, the field has made much progress in identifying the primate ventral visual pathway as crucial for housing neural circuits essential to object recognition^4,20,5^. A critical factor that led to progress in this domain has been the availability of rhesus macaques as an animal model that can mimic human object recognition behavior^21,22^. Given the ability to invasively probe finer-grain neural mechanisms in macaques^23,24^, studies have shown that a linear combination of image-driven population activity distributed across the macaque inferior temporal (IT) cortex (at the apex of the macaque ventral visual pathway) can sufficiently predict human object recognition behavioral error patterns on a battery of tasks^5,25^. Remarkably, these responses are typically recorded in monkeys who passively view the images without actively engaging in (or learning) the task -- suggesting that these representations are primarily bottom-up^5,25^ and task-independent^26^. Furthermore, a significant effort to model the transformations that follow the retinal responses (driven by the image) and culminate into the pattern of activity in IT has recently come in the form of a set of artificial neural networks (ANNs) that can partly explain the neural responses along these pathways^13,27,28^. Therefore, a reasonable approach to probe the mechanisms underlying the visual processing of scene context is to ask if macaques also mimic human context-driven behavior. If so, one could empirically probe the underlying neural mechanisms and compare current ANNs’ ability to explain those representations.

Interestingly, while current ANNs have been able to partially explain neural responses in V1^29^, V2^30^, V4^27,31,32^, and IT^27,33,34^, and many aspects of object recognition behavior^22^, recent studies have also shown that these models are heavily biased by the visual context during their training^13^ which lead to their misalignment with human behavior. These models also develop specific biases (e.g., shape-texture bias) that do not align with human strategies^35–37^. With the increasing evidence of discrepancies between ANNs and human behavior, it is critical to figure out how these models can be improved. The ability to probe context-dependent behavioral biases in monkeys and their underlying neural mechanisms allows us to develop strong constraints that can guide future model development.

In this study, we first developed quantitative behavioral metrics (coarse to fine-grained) to evaluate the psychophysical effects of contextual changes during object discrimination. We then conducted a thorough comparative analysis of the behavior of humans and monkeys. We further performed large-scale neural recordings across the macaque IT cortex to probe the strength of the image-driven IT responses and explain the observed behavioral variances. We contrasted the IT representations with those retrieved from the current most human-aligned ANNs. Our results unveil a nuanced understanding of how context influences object recognition in biological and artificial systems, which highlights significant parallels but also divergences in how humans, monkeys, and ANNs process visual context information.

## Results

We investigated the behavioral effects of scene context on humans and macaques during recognition of real-world objects, such as cars, animals, and fruits. We introduced multiple variations of the contextual information to further our understanding of what aspects of the object’s surrounding impact recognition. These variations include incongruent context, no context, and blurred context, among many others (**Fig 1A**). We developed a binary delayed match to sample object discrimination task (**Fig 1B**), where the participants, humans (**Fig 1C**) and monkeys (**Fig 1D**), identified the Target object shown in a sample test image (with varying contexts) when probed with two object choices (a target and a distractor). We quantified context-driven behavioral responses in both species with multiple quantitative metrics and assessed how well these metrics matched each other. Next, to probe the nature of the neural representations that could support these behavioral patterns, we examined how well the shared variance in their behavior is explainable by neural data from the inferior temporal (IT) cortex and the IT-like sub-units of current ANN models of primate vision (**Fig 1E**).

**Fig 1.**
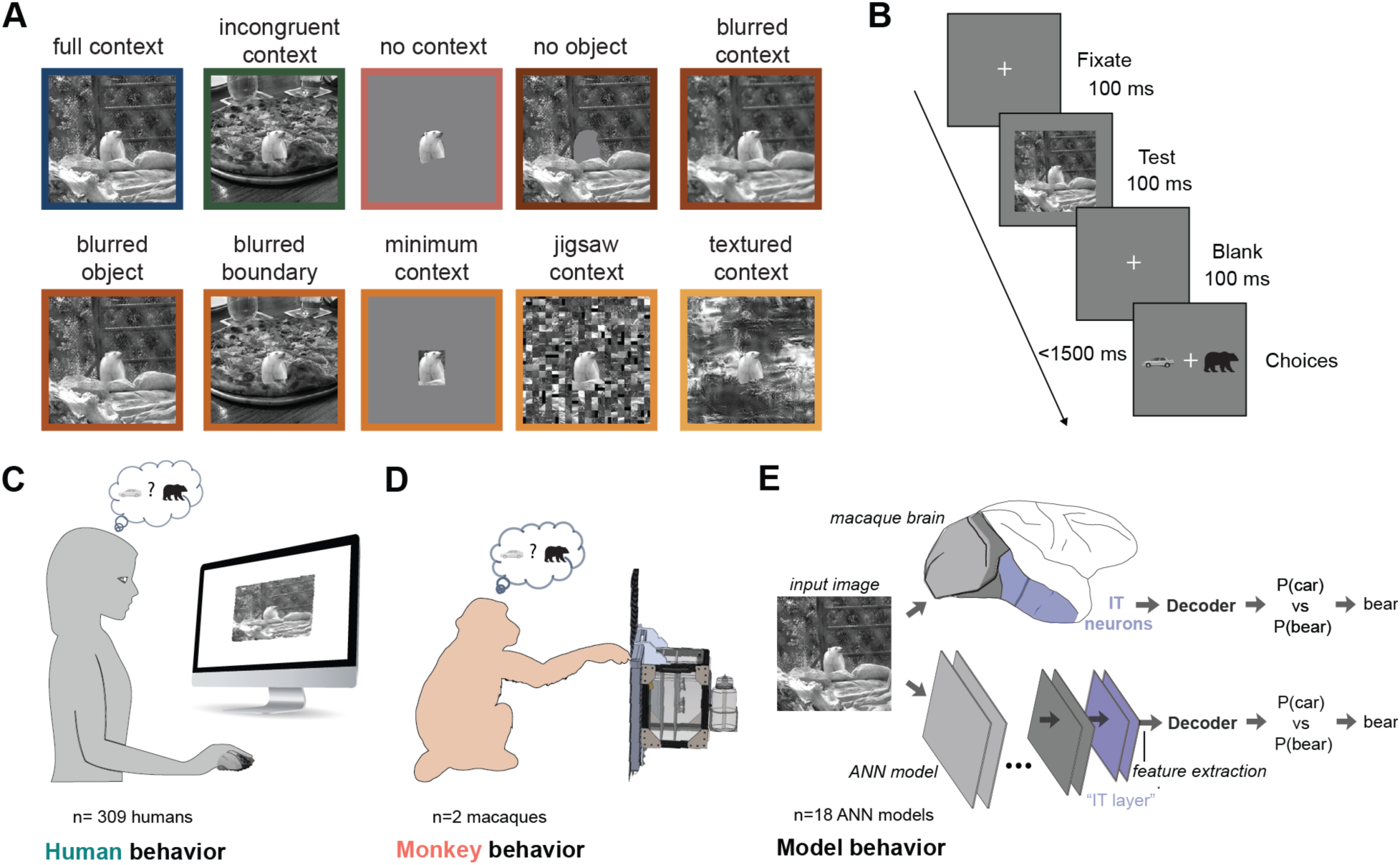
Comparing the influence of context in object discrimination performance across humans, monkeys and artificial neural networks (ANNs). **A**. Example of the ten contextual manipulations for one image of the set used for the experiments (details in Methods). The frames around each image indicate the color associated with that context type (only used for reference in the article, not in the actual experiments). **B.** Binary object discrimination task, showing the timeline of events for each trial. Subjects fixate on a cross, then the test image containing one of ten possible objects and contextual manipulations is shown at the center of the visual field (subtending 8 degrees of visual angle) for 100 ms. After a 100-ms delay, a canonical view of the target object (the same category as, but not a template match to, the test image) and a distractor object (one of the other nine objects) appears. The human or monkey indicates which object was present in the test image by clicking on one of the two choices. **C.** Schematic of the human behavioral task for 309 participants recruited from Amazon MTurk. **D.** Schematic of the monkey behavioral task for two context-trained adult macaques. **E.** Schematic of the model behavioral task for eighteen pre-trained ANN models (bottom, details in Table 1) and the neural data (top). To make the artificial models compatible with the specific primate binary object discrimination task, their most IT-similar feature representations were extracted and used to train the decoder - a multiclass SVM classifier - calculating the cross-validated probabilities for each object class in a one-vs-all manner. The model output is then the object class with the highest one-vs-all probability. Similarly, the most reliable neural responses (n=122 neural sites) from two context-naive monkeys were used to train the decoder and obtain the object class probabilities.

**Table 1.** Summary of the ANN models used grouped by training objective.

| Model | Architecture | Layer used |
| --- | --- | --- |
| **Image classification models trained on ImageNet** |  |  |
| AlexNet<sup>43</sup> | Generic CNN | features.12 |
| VGG-19<sup>44</sup> | Generic CNN | features.27 |
| MobileNet-v2<sup>45</sup> | Generic CNN | features.15 |
| ResNet-18<sup>46</sup> | Skip connections CNN | layer4.1 |
| ResNet-50<sup>46</sup> | Skip connections CNN | layer4.2 |
| ResNet-101<sup>46</sup> | Skip connections CNN | layer4.2 |
| ResNet-152<sup>46</sup> | Skip connections CNN | layer4.2 |
| DenseNet-201<sup>47</sup> | Skip connections CNN | features.transition3.pool |
| ConvNetXt Large<sup>48</sup> | Skip connections CNN | avgpool |
| GoogleNet<sup>49</sup> | Inception block CNN | inception5b |
| Inception-v3<sup>50</sup> | Inception block CNN | Mixed_7c |
| RegNetX 32GF<sup>51</sup> | Generic CNN | trunk_output.block3.block |
|  |  | 3-12.activation |
| ViT-b32<sup>52</sup> | Transformer | encoder.layers.encoder_layer_11.ln_2 |
| Swin-b<sup>53</sup> | Transformer | features.7.1.norm2 |
| **Image memorability models trained on LaMem** |  |  |
| MemNet<sup>54</sup> | Generic CNN | pool5 |
| ResMem<sup>55</sup> | Skip connections CNN | features.layer4.2 |
| **Object detection models trained on Microsoft COCO** |  |  |
| FasterRCNN (ResNet50 backbone)<sup>56</sup> | Skip connections RCNN | backbone.body.layer4.2 |
| RetinaNet (ResNet50 backbone)<sup>57</sup> | Skip connections RCNN | backbone.body.layer4.2 |

### Quantifying Context-Driven Changes in Object Recognition through Behavioral Metrics

To characterize how scene context influences the behavior of biological and artificial visual systems during object recognition, we developed quantitative metrics beyond the overall performance accuracy across all images. These metrics include the behavioral signature at the context level (B.C1, Behavioral, Context-Level 1-dimensional; see Methods) and a more fine-grained image level (B.I1, Behavioral, Image-Level 1-dimensional). The **context-level performance metric, B.C1** (human performances shown in **Fig 2A** - *right*), assesses the overall object discriminability within each context category (C). It does so by pooling accuracies across all images of a given context type (C) and all combinations of target and distractor pairs for those images (see Methods). This approach provides a broad understanding of how context influences recognition performance on a categorical level. In contrast, the image-level metric, B.I1 (detailed in Methods, human performance shown in **Fig 2A**, *left*), focuses on the discriminability of individual images, assessing how well the system distinguishes each object (O) from all others per image across varying contexts. This finer-grained metric allows for a more detailed analysis of performance variations at the image level. Expanding upon this foundation, we then seek to estimate the shared behavioral variance between humans and monkeys (behavioral signatures shown in **Fig S1A**), as depicted in **Fig 2B**. This comparative analysis could reveal one of the following scenarios. First, given species level differences^38^, we might observe that monkeys do not process visual context in the same way as humans and, therefore, exhibit no shared variance with humans (H0; **Fig 2B** - top panel). Second, it is possible that monkeys only share a fraction of variance with humans (H1; **Fig 2B** - middle panel). Lastly, it is also possible that within our set of tasks, images, and contextual variations – monkey and human behavior fully align with each other (H2; **Fig 2B** - lower panel). These conditions can be independently assessed for each of our behavioral metrics, and we expect that finer-grained metrics will enable us to more rigorously quantify the boundaries of the shared behavior between these two systems.

**Fig 2.**
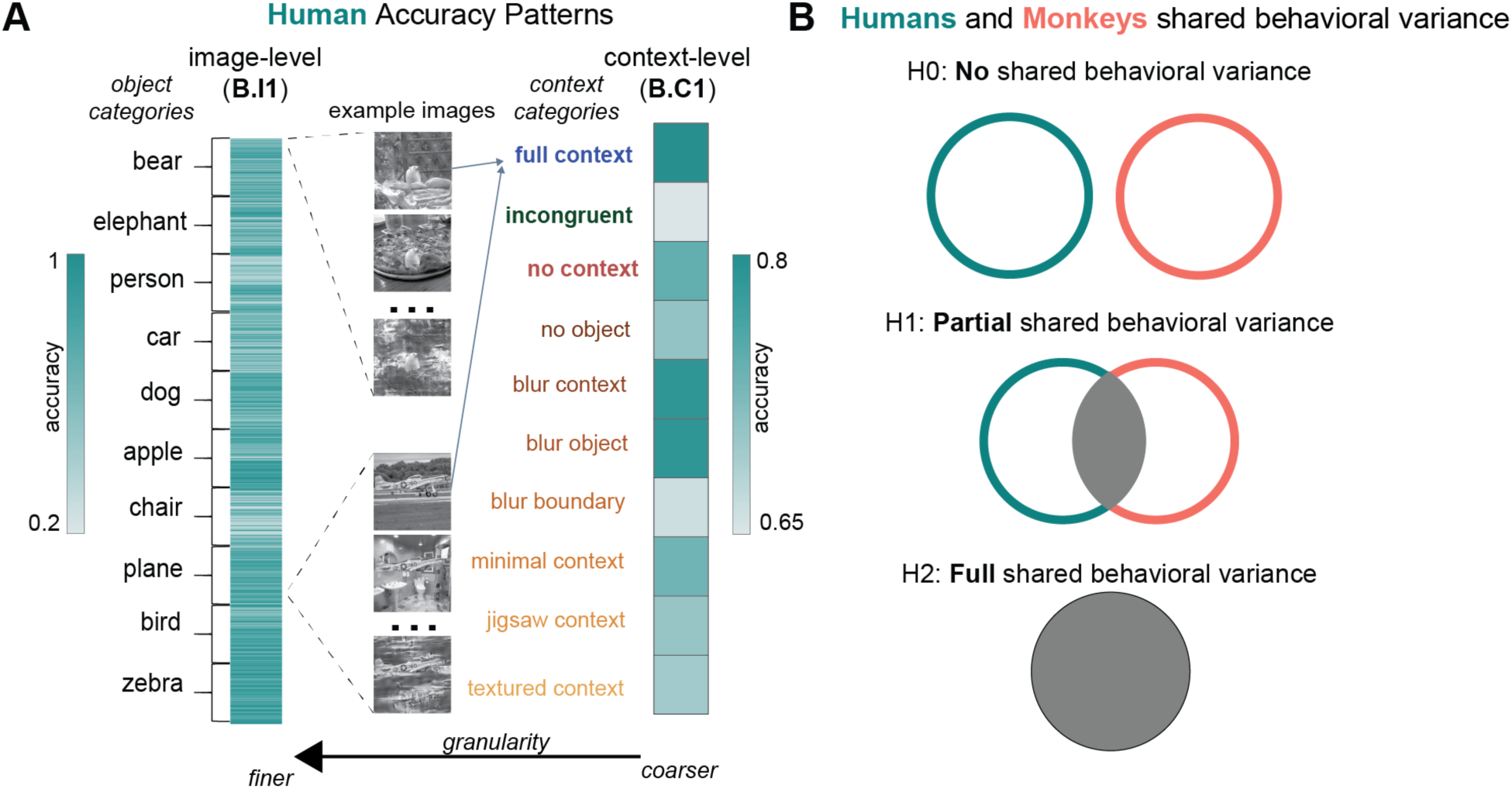
Behavioral metrics to quantify context-driven variations in task performance. **A.** Human accuracy patterns at an image-level (fine granularity, B.I1, left) and context-level (coarse granularity, B.C1, right). Each element of the B.I1 vector represents the overall accuracy (averaged across all tasks) for an image. A few example images are shown in the middle panel grouped by object category. The context-level signature (B.C1, right panel), is obtained by averaging the B.I1 values for all images of each specific context type (see **Fig 1** for examples of all the context types). The light and dark teal colors indicate lower and higher performances (see color scale next to each signature). **B.** The three hypotheses on the human-monkey shared behavioral variance.

### Object context induces significant changes in human behavior

Humans (309 participants on Amazon Mechanical Turk) participated in a binary object discrimination task (**Fig 1B**, for details, see Methods). Our results show that varying the context of the image changes the performance of the human participants. For instance, consistent with previous research^10,13,14^ humans show a significant reduction in accuracy for incongruent compared to congruent contexts (ΔAccuracy = 0.13 ± 0.21; Lilliefors test: full context p=0.004, incongruent context p=0.371, non-normal distribution, p>0.005; Wilcoxon rank-sum test statistic=4.4, p=0.0001; **Fig 3A**: blue vs green bars). The effect of contextual manipulations resulted in a consistent pattern of behavior (with a trial-split reliability of approximately 0.74, see **Fig S2A**, reliability across context types in **Fig S2B**). This high self-reliability is critical to ensure that contextual effects can be compared across animals, across species, and from biological systems to ANN models. The decline in accuracy for incongruent (compared to congruent) context was not solely due to the abrupt transition from the background to the object; even when the context/object boundary was blurred (termed blurred boundary), we observed the same effect. Predictably, removing the object, retaining only its silhouette, also led to reduced accuracy; however, performance remained well above chance, indicating that the context alone (with the object outline) provided enough information for accurate object discrimination. Moreover, when the context was removed or minimized, there was again a decrease in performance, confirming that humans also rely on the surroundings for object recognition. The blurring process itself seemed to have minimal influence on human responses, as the kernel size used was relatively small (see Methods). Using a synthesized texture (textured context), which retained the visual attributes of the original context, also adversely affected human behavior. Our results align with extensive research on context modulated human behavior^7,13^ and notably extend beyond the scope of previous work. In particular, we provide quantitative results from a forced binary choice task for a wider range of context variations. We define two behavioral signatures, allowing a coarse and fine-grain comparison within the human population and, importantly, across species - similarity with rhesus macaques. The cross-species consistency, coupled with access to the macaques’ neural circuits, provides a path for studying the neural mechanisms underlying contextual processing.

**Fig 3.**
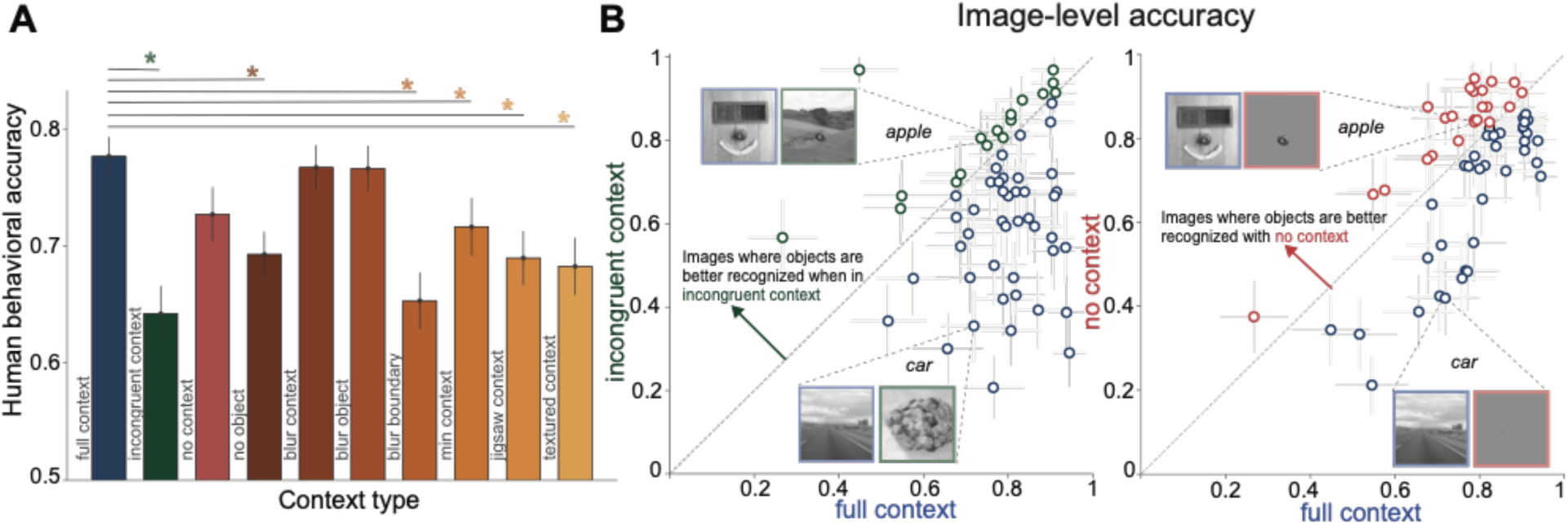
Context-driven changes in human behavioral task performance. **A.** Contextual manipulations produce significant changes in human visual recognition. Accuracy (mean 0.71±0.05) for each contextual manipulation (B.C1, **Fig 2**), with standard error across images. Statistics are shown for full context compared to other context variations (* denotes independent t-test, p<0.05). **B**. Left: Image-level accuracies (from the object discrimination task, with standard error across image trials) for full and incongruent context, each dot represents the human behavioral accuracy for the same object embedded in either full or incongruent context. Example images are shown where the object is better predicted in each context variation. Right: Similar as left, but comparing the accuracy for objects embedded in full context vs removing the context. Note that the car object is very small (< 1 degrees of visual angle) and hard to see without a lot of zoom (images are presented at the center of the visual field subtending 8 DOV angle, **Fig 1**).

### Object context induces significant changes in monkey behavior

To establish macaques as an appropriate animal model to probe the neural mechanisms of human context processing it is critical to first ask whether macaques behaviorally show similar contextual effects to humans. To ensure that macaques are familiar with the scene context per object category, we first explicitly trained them with images in full context (from the Microsoft COCO dataset, 160 images per object for ten objects). Macaques showed robust cross-validated accuracy during such training (**Fig S1B**).

Once the monkeys (n=2) were fully trained (i.e., reached ≥80% performance) in their home cages (see learning curve **Fig S1B**), we measured their object discrimination performances with the same contextually manipulated images as humans (**Fig 1A**). Monkey behaviors were highly reliable (as measured by trial split-half reliability, r=0.76, see Methods, **Fig S3A**), and correlated with each other at both the context level (corrected Pearson R=0.98, corrected by both monkeys’ self-consistency, see Methods, **Fig S4A**), and at the image-level (corrected Pearson R=0.83, **Fig S4B**). Similar to humans, monkeys also showed a significant reduction in accuracy for incongruent compared to congruent contexts (ΔAccuracy = 0.104 ± 0.18; Lilliefors test: full context p=0.173, incongruent context p=0.58, normal distribution, p>0.005; independent t-test, t(59) = 3.305, p=0.001, **Fig 4A**: blue vs green bars). **Fig 4B** compares the trial averaged image by image accuracy between full and incongruent context (left), as well as full and no context (right). At the individual image level, we observe some images for which the object placed in an incongruent context was better recognised than when the same object was embedded in a congruent context (see example of an apple in **Fig 4B**, left). Similarly, some objects were better recognized when fully removing the context compared to keeping the full congruent context (see apple example in **Fig 4B** right).

**Fig 4.**
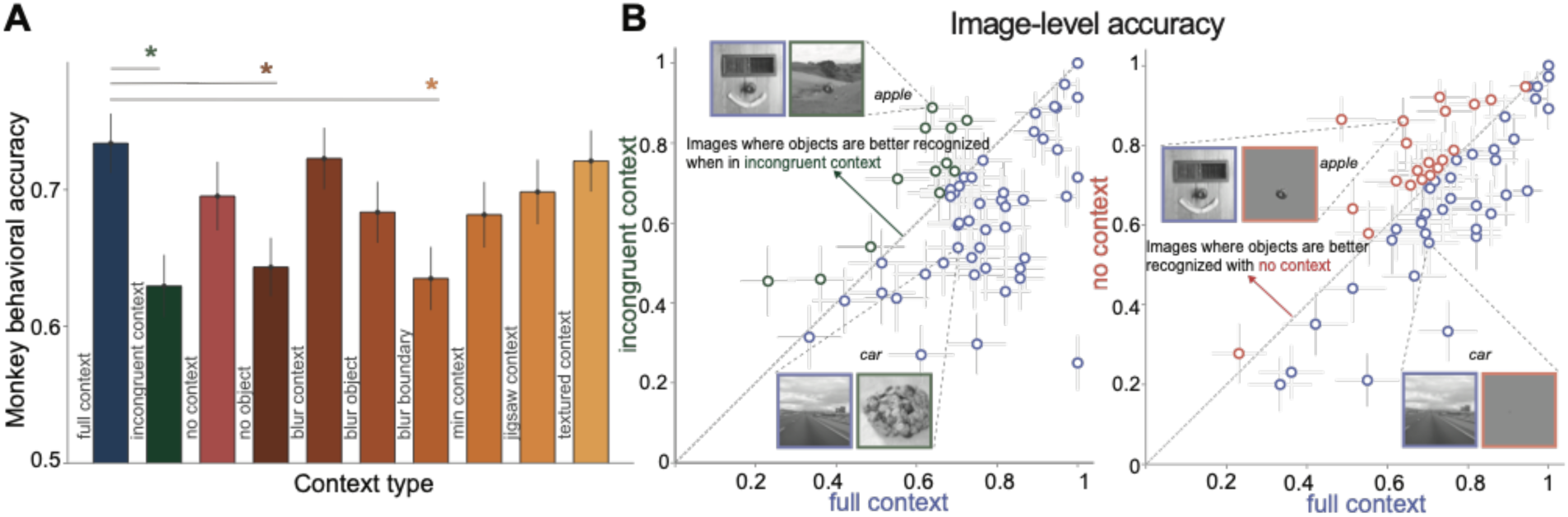
Context-driven changes in monkey behavioral task performance. **A.** Contextual manipulations produce significant changes in monkey behavior. Accuracy (mean 0.68±0.04) for each contextual manipulation (B.C1), with standard error across images. As in **Fig 3A**, statistics are shown for full context compared to other context variations (* denotes independent t-test, p<0.05). Chance = 0.5. **B.** Left: Image-level accuracy for incongruent context versus full context (format as in **Fig 3B**), each dot represents the pooled monkeys’ behavioral accuracy for the same object embedded in incongruent (y-axis) or full (x-axis) context with standard error across image trials. We show example images where the object is better recognized in each context variation. Right: Similar as left, but for no context (y-axis) versus full context (x-axis).

### Humans and monkeys share significant variance in context driven changes in object recognition

We directly compared monkey and human performance for the same images and task. Our results show a remarkable consistency between monkeys and humans at the context level (C1 corrected Pearson R=0.83, **Fig 5A**). For example, both monkeys and humans performed best in the full context condition (blue point) and worst in the incongruent context condition (green point). However, the majority of points in **Fig 5A** fall below the diagonal, indicating that humans outperformed monkeys in most context conditions (Δ (human - monkey) =0.02±0.03, Lilliefors p=(0.517, 0.487) : normal distribution; paired t-test, t(9) = 1.86, p=0.1). The two exceptions were the jigsaw and textured context conditions, where monkeys slightly outperformed humans. To quantify the variability across humans, we calculated the human ceiling by comparing the shared variance between two separate pools of human subjects (teal band in **Fig 5B**). We then compared the shared human-monkey variance to this human ceiling. Since we are comparing a pooled population of 309 humans to the n=2 monkey pool, we looked at the effects of monkey pool size on its consistency with human data. As the number of monkeys in the pool increased from one to two, the shared human-monkey variance increased by 4.3% (gray bars in **Fig 5B**). Extrapolating to an infinite pool of monkeys using a “pseudo” human consistency function (sigmoid) derived from subsampling the human pool, we estimate that the asymptotic shared variance between monkeys and humans would reach approximately 80% of the human ceiling. Next, we compared monkey and human performance at the individual image level (**Fig 5C**). Again, we found a significant correlation between monkeys and humans (I1 corrected Pearson R=0.63), although the relationship was weaker than at the context level. The slope of the regression line in **Fig 5C** suggests that humans outperformed monkeys on average, but this difference was not as pronounced as at the context level (Δ (human - monkey) =0.02±0.18, Lilliefors p=(0.001, 0.001): non-normal distribution; Wilcoxon test: statistic = 79913.5, p=0.02). The shared variance analysis at the image level (**Fig 5D**) revealed that humans were less consistent with each other compared to the context level (**Fig 5B**), as expected due to the increased granularity of individual images. This effect was even more pronounced for monkeys, with a larger drop in shared variance at the image level compared to the context level. Increasing the number of monkeys in the pool from one to two improved the shared human-monkey variance by 8.2% at the image level (**Fig 5D**). Extrapolating to an infinite pool of monkeys, we estimate that the asymptotic shared variance would reach approximately 70% of the human ceiling at the image level.

**Fig 5.**
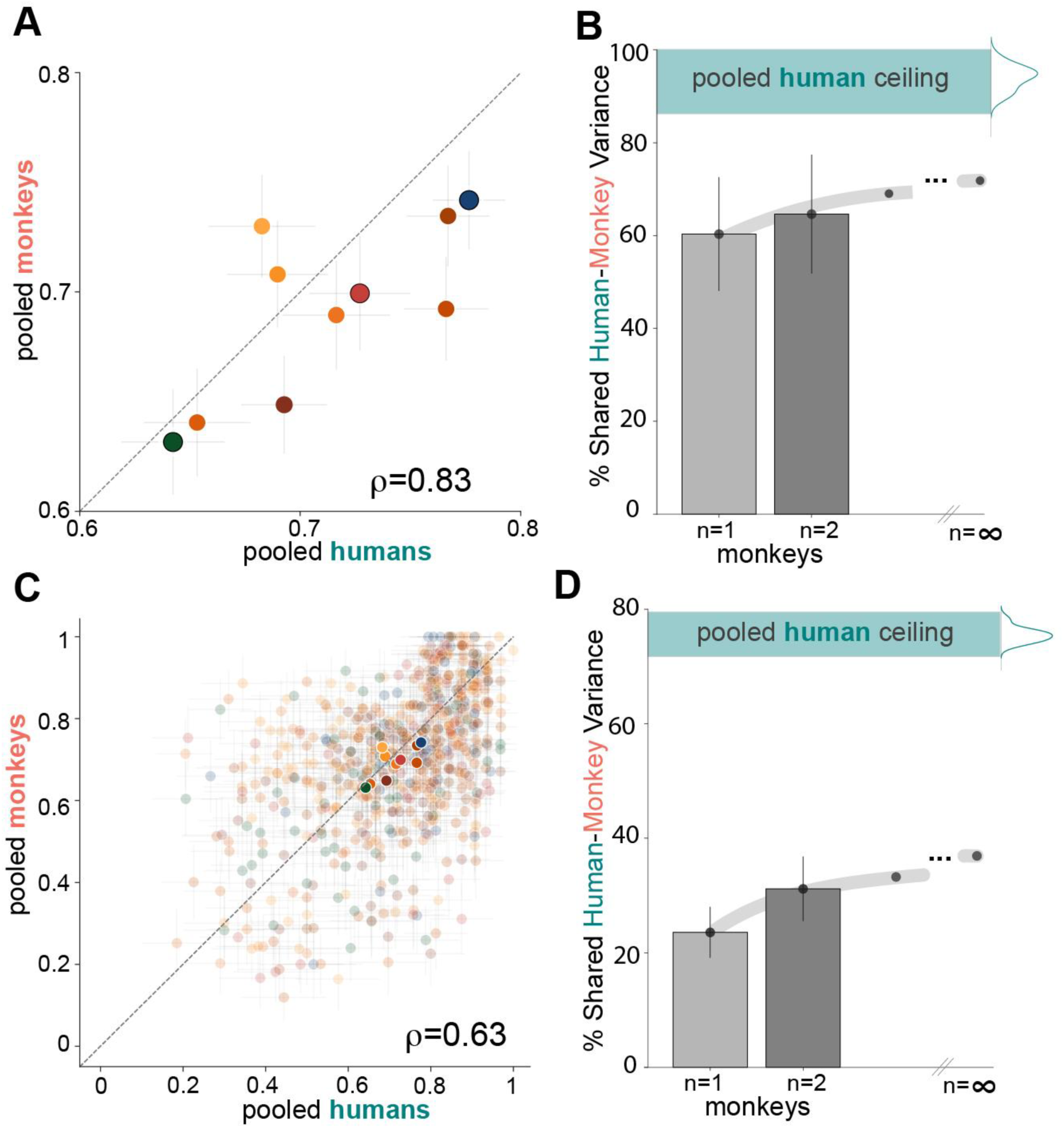
Monkeys and humans show similar (but not identical) context-driven behavioral changes. **A**. Context-level (B.C1) correlation between the pooled monkeys (n=2) and pooled humans (n=309). Each point represents the mean accuracy for a contextual variation with standard error across images of that context type (colors as in **Fig 1A**, pooled monkeys mean 0.69±0.20, humans mean 0.71±0.18). The three main context types: full (blue), incongruent (green) and no context (red), are shown with a black stride around the filled point. The value ρ indicates the noise corrected correlation coefficient (Pearson R). **B.** Shared human-monkey explained variance at a context-level (mean with standard deviation across context types), as a function of the number of monkeys used for pooling. The asymptotic value for an infinite pool of monkeys is obtained by extrapolating the “pseudo” human consistency function (Methods). The human self-consistency ceiling is shown as a teal band. **E.** Image-level correlation (B.I1) for the pooled monkeys and humans, each low opacity point shows the performance (mean accuracy) for an image with standard error over image trials, the higher opacity points are the B.C1 mean (from A), colors map to context types as defined in **Fig 1A**. **F.** Similar to C but mean shared variance at an image-level with standard deviation across image subsamples.

### Population activity across the IT cortex in a context-naive monkey fully explains the shared behavioral variance between humans and monkeys at the overall context-level

Our behavioral results (**Fig 5**) demonstrate that humans and macaques share a significant proportion of variance (context level shared EV=62.43%) induced by context variations during object discrimination. To understand the neural mechanisms behind these contextual influences, we require a more detailed examination of the neural networks involved. Previous studies have shown that IT population responses in monkeys (passively viewing images, see Methods) can be linearly combined to sufficiently explain human object category (and category-orthogonal) based behavioral patterns ^1,5,25^. Therefore, we aimed to assess the extent to which the image-driven responses in the IT cortex of context-naive macaques could account for the variance observed between humans and monkeys. Similar to the expected observations while comparing human and monkey behavior (**Fig 2B**), we hypothesized that there could be no overlap (*H0*; **Fig 6A**), partial overlap (*H1*; **Fig 6A**), or full overlap (*H2*; **Fig 6A**) between the neural predictions and primate behavior.

**Fig 6.**
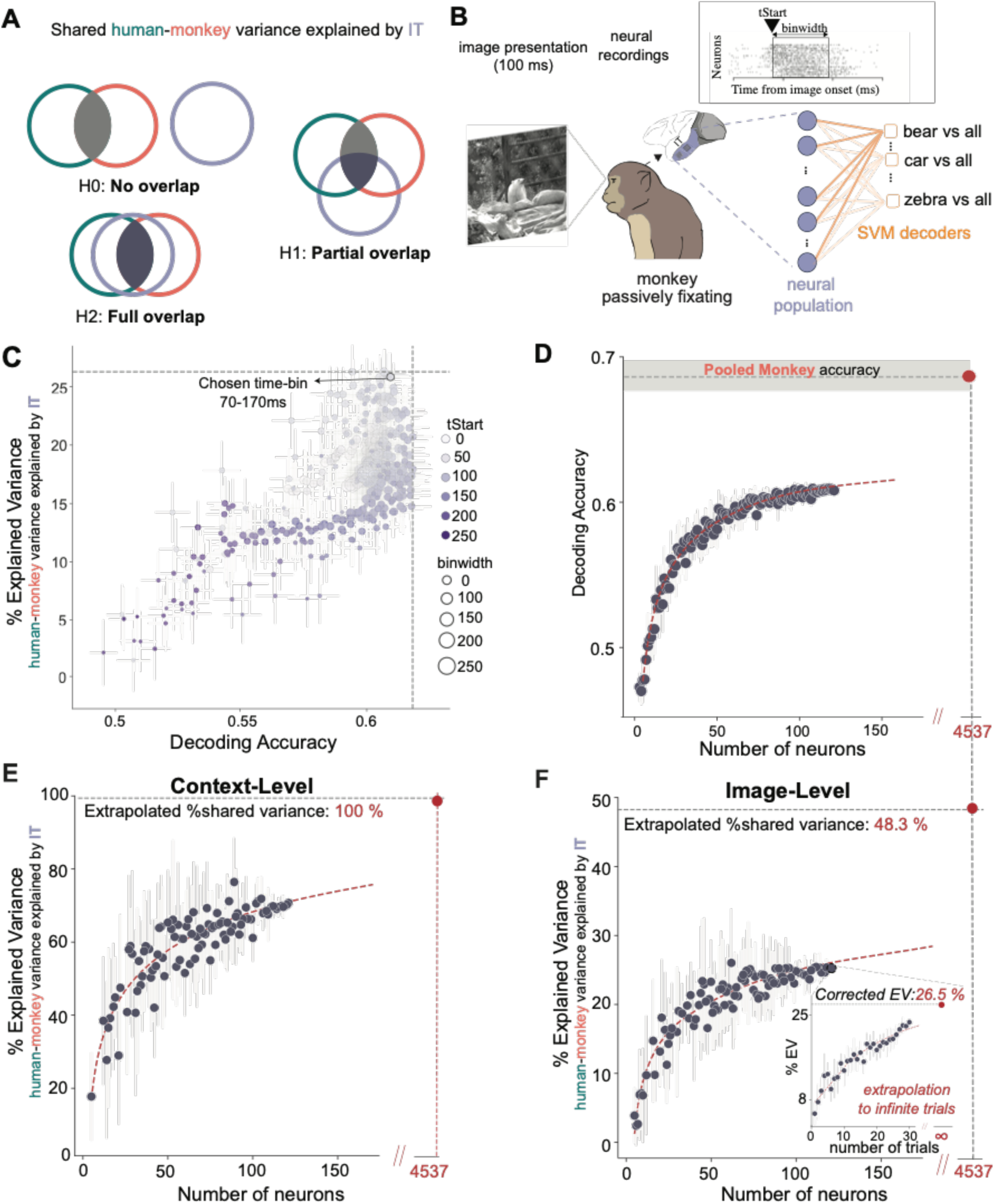
Context naive macaque IT fully explains the human-monkey shared behavioral variance at a context-level but only partially at an image-level. **A**. The hypotheses for how much of the human-monkey shared explained variance (HM-EV) can be explained by IT. **B.** The neural data was recorded while the monkey was passively fixating on the center of an image (8 degrees of visual angle) presented at the animal’s center of gaze for 100 ms. The object category decoding was done by training a multi-class SVM classifier (one vs all for each object category) tested in a cross-validated way on the same images and tasks as those presented to humans and monkeys. For a given image, the decoding output is the object class with the highest one vs all probability. All behavioral predictions from the decoder were for images where the object was not seen in any phase of the model training, making sure we never show an image of the same object (regardless of the contextual manipulation) during the fitting and testing. We decoded the object category from each possible time-bin of the neural data by varying the tStart (start of the time-bin with respect to image onset, in ms) and binwidth (length of the time-bin, in ms) of the obtained neural population vector (0-300 ms per image presentation). **C.** Results from decoding all time-bins (filtered with self-consistency >0.1) from the neural data, color indicates the bin start, size indicates the bin length. The percent of image-level explained variance from the shared human-monkey variance is shown (y-axis, with standard error across image subsamples) as a function of the decoding accuracy for each bin (x-axis, with standard error across images). We used the 70-170 ms time-bin for all subsequent analyses. **D.** Decoding accuracy with standard deviation (one-vs-all accuracy, chance level = 0.5) across neuron subsamples for the 70-170 ms time-bin, as a function of the number of neurons . An extrapolation (dashed red curve) estimates the decoded accuracy from a neural population of 4537 recorded neural sites would reach the overall pooled monkey accuracy (0.69, gray band shows monkey accuracy mean with standard error across the 600 images). **E**. Context-level variance explained by the neural data, from the HM-EV. The EV is obtained by subtracting the HM-EV when controlling for the neural data (partial correlation) from the full HM-EV and normalizing by the full HM-EV (see <u>Methods</u>). We show the EV as a function of the number of neurons used for decoding, showing an extrapolation to 4537 neurons would fully explain the B.C1 HM-EV. Each point shows an average (with standard deviation error bar) across ten different subsamples of neurons used, corrected by extrapolating to an infinite number of trials for those specific neurons. **F.** Similar to E, but for image-level shared variance. The inset shows the correction for the EV for 122 neurons by extrapolating to an infinite number of trials as done for context-level, each point shows an average (with standard deviation error bar) across ten different subsamples of trials. The extrapolation of the EV to the number of neurons needed to reach monkey accuracy (see decoding accuracy extrapolation in D) gives a ceiling of 48.3% of image-level human-monkey behavioral variance that can be explained by the context naive monkey IT neural data.

We performed chronic neural recordings using Utah arrays across the IT cortex in two macaques that passively viewed the images (used in the behavioral tasks) presented for 100 ms each (**Fig 6B**, see Methods). We combined the most reliable neural sites (n=122; see criteria in Methods, 30 sites from monkey 1, 92 sites from monkey 2) across the two monkeys to generate a pooled neural population for further analysis. Similar to previous methods^5,23,28^, we used linear classification-based algorithms (**Fig 6B**) to decode the object category for each image from the pooled neural data and estimated the neural predictions for the behavioral metrics (explained above, e.g. C1, I1).

We first asked how well the macaque neural responses can predict the shared variance between humans and macaques at the B.C1 level. Therefore, we performed a partial correlation analysis between human and macaque C1 behavioral patterns while controlling for the IT population activity-based predictions of B.C1. To account for the irreducible noise in the neural data, we corrected the partial correlation by extrapolating it to an infinite number of trials for the neural data (see inset **Fig 6F**). Interestingly, the neural data (122 sites) explained 75% of the context-level shared monkey-human C1 variance (**Fig 6E**). To further address the data limitations arising from the limited number of neural recordings, we extrapolated the neural decoding accuracy to match the monkey accuracy (logarithmic function, **Fig 6D**). This extrapolation led to an estimation of 4357 neural sites needed to reach monkey accuracy. A logarithmic extrapolation of the explained variance (EV) to 4357 neural sites indicates that IT would fully explain the human-monkey B.C1 variance if we had more neural recordings (**Fig 6C**).

### Population activity across the IT cortex in a context-naive monkey only partially explains the shared context-driven behavioral variance between humans and monkeys at the image-level

To further stress test whether IT responses from untrained (task-naive) monkeys can explain finer grained behavioral patterns, we next turned to predictions for the I1 level (image-level shared variance). As shown in **Fig 6F**, the recorded reliable neural population (122 neural sites) explains only a fraction of the image-by-image behavioral variance (up to 25%). This result suggests that the context-naive IT population may not capture all the necessary information to fully predict the shared human-monkey behavioral patterns at the image level. To address the possibility that the limited explained variance might be due to the restricted number of recorded neurons, we applied the same extrapolation method as used for the context-level EV (**Fig 6E**). We estimated that approximately 4357 neural sites would be needed to match the pooled monkey behavioral accuracy (**Fig 6D**). However, despite this extrapolation, the neural data from context-naive IT could not fully explain the image-by-image shared primate variance, reaching a ceiling of only 48.3% (**Fig 6F**). The discrepancy between the context-level and image-level explained variance highlights the complexity of the neural mechanisms underlying context-dependent object recognition and the limitations of using context-naive neural responses to predict fine-grained behavioral patterns. In summary, while the context-naive IT population activity can fully explain the shared human-monkey behavioral variance at the context level (**Fig 6E**), it only partially accounts for the variance at the image level (**Fig 6F**). This finding underscores the need for further investigation into the neural mechanisms that shape the shared behavioral patterns between humans and monkeys in the presence of contextual variations.

### Low-level image-based features do not explain the shared human-monkey behavioral variance

So far, we have observed that human and macaques share a significant amount of behavioral variance both at the coarse (B.C1) level and the finer-grained (B.I1) level. The image-driven task naive IT responses can fully explain the C1 variance but not the I1 level variance. We next asked how much of these results can be explained by low-level image features. For every image, we extracted a range of basic image features, such as object size, location, and category, spectral mean and standard deviation(std), and contrast mean and standard deviation (std) (**Fig 7A**). The low-level features were chosen to capture basic properties of the images that could potentially influence object recognition performance, such as the object’s saliency (contrast) and its placement within the scene (location). We observed that these low-level features do not explain the context-level (**Fig 7B**, 55.39±7.08% mean±95% CI of noise ceiling, max low-level feature EV = 3.03±2.2%) or image-level (**Fig 7C**, 23.4±2.9% mean±95% CI of noise ceiling, max low-level feature EV = 2.47±1.47%) measured shared behavioral variance. Among all the low-level features tested, object size showed the most consistency with the shared human-monkey variance at the image level, aligning with prior studies highlighting its influence on human behavior (Zhang et al., 2020). In particular, the positive correlation with human and monkey performance was more significant for smaller object sizes and diminished for objects with size beyond 5 degrees of visual angle (**Fig S5**). Its effect, however, is marginal, accounting for only 10% of the shared human-monkey image-level variance. This suggests that while object size plays a role in shaping the shared behavioral patterns, it alone cannot fully explain the observed consistency between humans and monkeys. Taken together, we infer that low-level image features alone are insufficient to explain the shared behavioral patterns observed in humans and monkeys, indicating the need for more complex, higher-order processing to fully account for the context-dependent object recognition performance.

**Fig 7.**
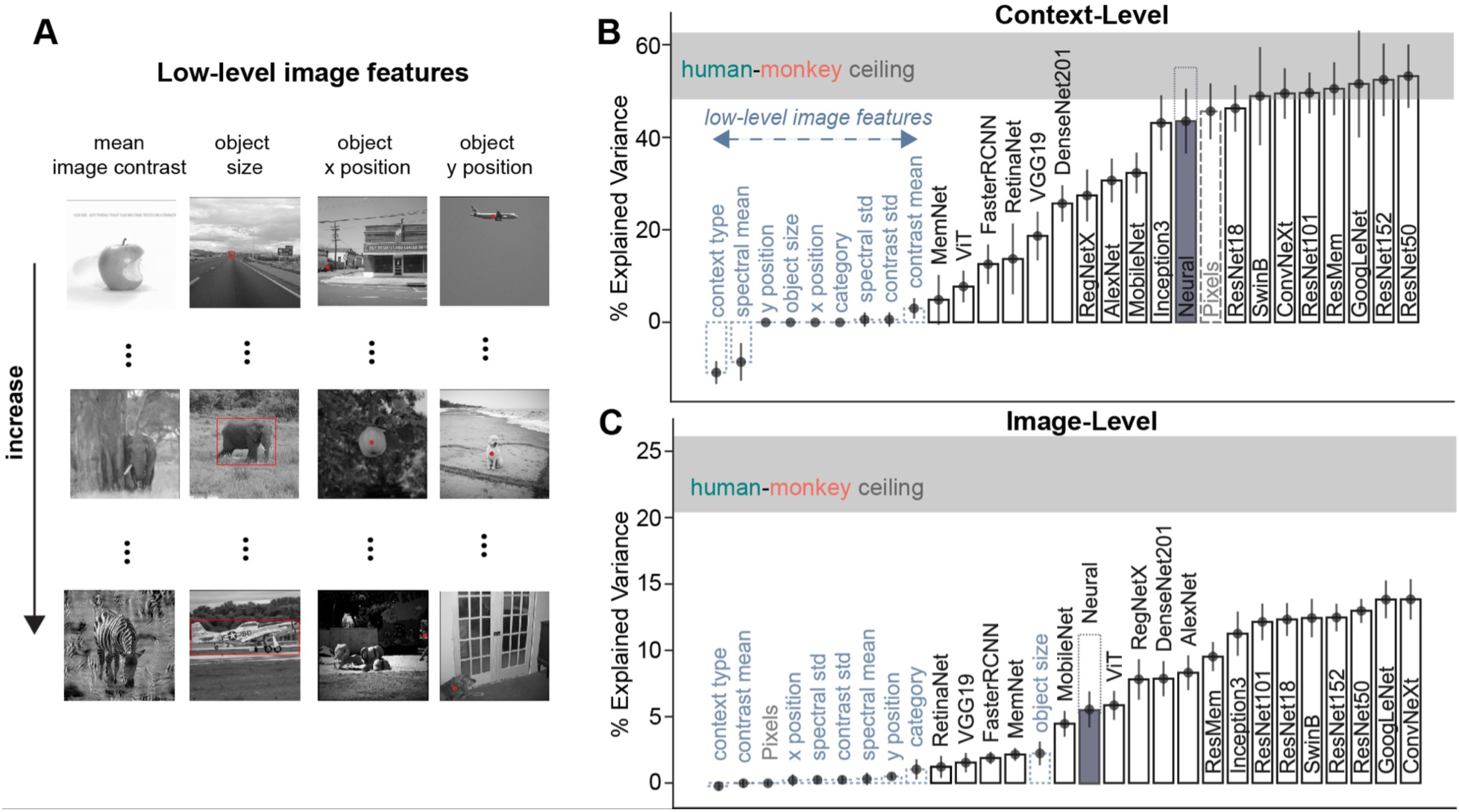
ANNs fully explain the human-monkey shared behavioral variance at a context-level but only partially at an image-level. **A**. Example of images with increasing “intensity” (top to bottom) of 4 example low-level image features: mean image contrast, object size, object x and y position. The objects are noted with a red dot or bounding box. **B**. The human-monkey shared variance explained by the low-level features and ANN models at a context level, showing the mean fraction of explained shared variance with standard deviation across different image subsamples from 20 bootstraps (choosing 600 images with repetition). The low-level features are shown with dotted light blue bars. The ANN decoding was done in the same way as for the neural population (multiclass SVM), only using the extracted model features from the ‘IT’ layer for each model. Pixels’ (control model - flattened image pixels) performance is shown in a dashed gray bar. The mean human-monkey shared variance ceiling is shown in gray, with standard deviation across different image subsamples from 20 bootstraps (same as for the bars). We are noting the neural corrected EV (purple) when using all the recorded reliable neural responses (122), and the extrapolation (from **Fig 6**) with the dotted bar. **C.** Similar to B but showing the explained variance at an image-level.

### ANNs fully explain the overall context-level human-monkey shared behavioral variance

Next, we tested whether the current best models of primate vision, a family of deep convolutional neural networks (DCNNs), vision transformers, or recurrent convolutional neural networks, can predict the behavioral variance observed on a context and image-by-image level. Using a multiclass SVM decoder, we mimicked the same object discrimination task presented to humans and monkeys. We used the most ‘IT-like’ layer features from each model (for details, see Table 1), projected to a 3k lower dimensional space (via a Random Gaussian Projection). These model accuracy decodes showed sensitivity to contextual changes, with their accuracy varying across different context types and images, as shown in their behavioral signatures (**Fig S9**). The ANN models’ ability to capture context-dependent performance variations suggests that they have learned to extract and process contextual information in a manner that is relevant for primate object recognition.Similarly, for the neural data, we did a partial correlation analysis for the pooled monkey and human population behavioral patterns while controlling for each of the artificial models’ variance. Our results show that most models, including the Pixels control model, can explain the context-level shared behavioral accuracy patterns of humans and monkeys.

### ANNs only partially explain the image-level human-monkey shared behavioral variance

While ANNs fully capture the shared human-monkey behavioral variance at the context-level, their performance at the finer-grained, image-level is less comprehensive. Despite the models’ ability to explain the overall context-dependent behavioral patterns, they struggle to account for the more intricate, image-specific variations in primate behavior. At B.I1 level, the models explain at most 70% of the image-level shared primate behavioral variance. This discrepancy is due to the finer grain accuracy variations within both context and object category types that are not consistently aligned between the primates and the artificial models. We found a strong correlation between the fraction of explained shared human-monkey variance and the decoding accuracy from the model features (Pearson R=0.78 for B.C1 and 0.95 for B.I1, Fig S7), indicating that improving the model accuracy could allow them to fully explain the shared human-monkey B.I1 behavioral variance. The control Pixels model - using the raw image pixel values, capturing the context-level shared behavioral patterns, was falsified at an image level. This reveals more complex image-level shared behavioral patterns that are not due to the raw image features. The image-level gap was consistent when comparing at an individual level - these models could not fully explain the (full) human, monkey or neural image-level behavioral patterns (**Fig S2**, **S3** and **S8**). This indicates that such models do not currently possess the mechanisms required to process scene context in a primate-like fashion.

In summary, while low-level image features and current artificial neural networks can account for the overall context-level shared behavioral variance between humans and monkeys, they fall short in fully explaining the more intricate, image-level behavioral patterns. These findings highlight the need for further advancements in artificial neural network architectures and training paradigms to better capture the nuanced, context-dependent object recognition processes observed in primates.

## Discussion

In this study, we highlighted the critical role of context in primate object recognition. The visual object recognition abilities of monkeys that were initially trained to categorize objects in their natural context were strongly modulated when we deliberately varied the contextual cues. Our findings reveal that both humans and monkeys exhibit a significant sensitivity to contextual cues, which goes beyond low-level image attributes. Indeed, macaques shared a significant variance in their context-driven behavioral error patterns with humans. Thus, we established rhesus macaques as a viable animal model for investigating scene context in human visual recognition, paving the way for further studies into the neural underpinnings of contextual modulation. However, our analysis also revealed that at the image-level, monkeys do not entirely mimic human behavioral patterns, suggesting potential limitations in the depth and duration of their training or inherent species-level differences in sensory processing and cognition. In addition, we observed that the population activity distributed across the IT cortex of naive monkeys that were not explicitly trained with objects in context do not fully explain the context-driven behavioral patterns of the context-trained monkeys. Furthermore, our ANN-based simulations further reveal the substantial impact of context on the predictive behavior of current ANN models. Notably, ANNs exhibit limitations in their explanatory power for image-level comparisons with primates under varying contexts, indicating a clear need for model enhancements to accurately mimic the complex influence of context in primate visual recognition.

### Context modulates visual object recognition in humans and monkeys

Our results underscore the critical role of context in primate object recognition, aligning with an extensive corpus of literature on visual cognition ^11,13,39^. This research has established that human visual object recognition capabilities are modulated by contextual cues. Such cues are informed by our understanding of object occurrence statistics, which dictate notions of congruency or incongruency within a given scene. Interestingly, our findings reveal that monkeys, much like humans, exhibit sensitivity to these statistical cues. Across diverse context manipulations, we observed substantial decrements in object discrimination accuracy compared to fully congruent scenes - up to 13% in humans and 10.4% in monkeys for incongruent contexts. These striking parallels between the two primate species underscore the viability of macaques as a model system for probing the neural computations underlying context processing. Critically, our results extend beyond prior work by demonstrating context sensitivity across a broad range of manipulations and employing rigorous, multi-faceted behavioral metrics designed to quantify performance changes induced by contextual cues. The tight concordance points to potential shared cognitive mechanisms, such as knowledge of object co-occurrence statistics, relative sizes, and positional regularities, which could account for the context facilitation effects observed in both species.

### Goodness of monkeys as a model of human contextual processing

While the largely consistent effects at the coarser, context-level, validate macaques as a model, some discrepancies remain. Our analysis revealed that at the image-level, monkeys do not entirely mimic human behavioral patterns. To familiarize them with various contexts—such as cars on roads, bears in the jungle, and chairs in rooms—we trained these monkeys extensively with natural photographs (from the MS COCO image dataset) until their performance plateaued, as shown in **Fig S1B**. Despite reaching high performance, the mismatch between human and monkey responses suggests that the depth and duration of training might not have been sufficient. Enhancing the training regimen could potentially lead to a better alignment with human context-level behavior, reducing the disparity observed in image-level variance. However, potential confounds like the limited stimulus set size and specific task demands cannot be ruled out either. Importantly, instances where context manipulations like blurring had relatively small impact on performance in both species provide insights into boundary conditions that inform and constrain models of contextual reasoning. Another critical consideration is the inherent species-level idiosyncrasies and differences in brain structures between humans and monkeys ^38^. These biological distinctions might inherently limit the degree to which monkeys can model human contextual processing. While further training might narrow the behavioral gap, some level of divergence might always persist due to fundamental differences in sensory processing, visual experience, and cognition between the two species. Understanding and acknowledging these limitations is vital as we continue to refine monkeys as models for human visual processing. Future research should explore both the potential and the boundaries of this animal model, aiming to optimize training strategies and deepen our understanding of the species-specific factors that influence contextual processing. Through this nuanced approach, we can better leverage the strengths of monkeys as models while being mindful of their inherent limitations.

### Insufficiency of ANN models to explain primate context-driven behavior

Deep ANNs are currently the best models of human vision and also show remarkable performance in computer vision tasks ^3,28^. These models have been trained extensively on images of objects in context from large datasets (typically ImageNet). Our findings show that while these ANNs were able to fully explain the context level (B.C1; the coarser metric) shared primate variance, they failed to completely capture the finer grain image level accuracy patterns (B.I1). Even simple pixel-based models could predict the broad variations in B.C1 (**Fig 7**), underscoring the limitation of such coarse metrics in capturing the nuanced differences in visual context processing. However, a shift in focus to finer, image-by-image level variations revealed a more intricate picture. At this granular level, we discerned the primary distinctions between humans, monkeys, and ANNs. While monkeys show partial overlap with human behavior, a significant portion of this image-level variance remains unexplained by current ANN models. This gap highlights a critical area where artificial systems diverge from natural primate visual processing, suggesting that while ANNs can mimic some aspects of primate vision, they still lack certain mechanisms that drive the nuanced, context-driven behaviors observed in humans and monkeys. These observations not only challenge the sufficiency of broad behavioral metrics in capturing the essence of visual context processing but also point to image-level analyses as a more sensitive and discriminating tool for understanding the subtleties of primate vision. The partial alignment yet notable divergence of current ANN models from primates points towards key computational mechanisms underlying context integration during object recognition that may still be lacking in artificial systems. Aspects like rapid integration of segmented objects with contextual associations and scene statistics, combination of high-resolution foveal and low-resolution peripheral representations, oculomotor sampling routines tuned for context (however, see^13, 14^), or other dynamic processes could be critical for human-level contextual reasoning. Pinpointing and distilling such mechanisms from the primate brain represent exciting future directions. We tested a range of models (Table 1), to gain further insight into the model architectures that could explain the B.I1 primate shared behavioral patterns better. We observed that deep ANNs with residual connections, as well as inception modules, are most aligned with human-monkey behavior (**Fig S6**). This indicates that allowing for feed-forward long-range dependencies between features (e.g., low-level features like edges with higher-level features) and preserving the finer-grained information from earlier layers (which can be lost due to the depth of models) by using bypass connections could benefit the alignment of these ANNs with primate behavior. Furthermore, ANN decoding accuracy (signatures in **Fig S9**) predicts the fraction of explained monkey-human shared variance (**Fig S7**), indicating that by improving the model’s decoding accuracy, we could come closer to bridging the I1 explainability gap.

### Role of IT cortex in processing scene context

The inferior temporal (IT) cortex is integral to visual object processing ^4,5,23,40^, yet our findings indicate that responses from context-naive monkeys may not fully encapsulate the representation of scene context akin to that in humans or context-trained monkeys. This shortfall calls for a nuanced approach in future investigations into the IT cortex’s role in context processing. One explanation for this is that our data might be sample-limited, affecting the breadth and depth of our inferences. Constraints such as the extent of IT neural data sampling, the diversity of images, trials, objects, and context variations might have curtailed our ability to fully capture IT’s capabilities in context processing. To address these limitations, we conducted extrapolation analyses (**Fig 6**) to estimate the scaling laws governing our data, aiming to predict how increasing our sample might influence our findings – further corroborating the insufficiency of naive IT-based decodes to explain human behavior. Secondly, the lack of refined representational capacity in the IT cortex of naive monkeys might be due to insufficient exposure to varied contextual cues – improving which might amplify the IT cortex’s ability to represent scene context. Additionally, investigating the interaction of the IT cortex with other brain regions, both within the ventral stream like areas V4 and outside the ventral pathway such as the ventrolateral prefrontal cortex (vlPFC), and their correlation with behavior in trained and untrained monkeys could illuminate new aspects of neural processing. This exploration is crucial to discern whether other areas might compensate for or augment the IT cortex’s function in context processing, thus providing a more holistic view of the neural networks at play in this intricate task. Together, these strategies will deepen our understanding of the IT cortex’s role and pave the way for a more comprehensive grasp of the neural underpinnings of context processing in vision.

By bridging behavioral, computational and neural levels of analyses^41^, we can develop integrated accounts reconciling the cognitive influences of context with their neural underpinnings and use them to inspire more neurally-grounded computational models. Overall, this multi-pronged approach paves the way for a deeper understanding of how context facilitates robust object perception across primates.

## Methods

### Visual Stimuli

We generated an imageset comprising 600 grayscale images from 10 object categories (bear, elephant, person, car, dog, apple, chair, plane, bird, zebra). For each object category, we selected six natural images from the Microsoft Common Objects in Context (COCO) dataset, varying in object size and location, which were center cropped, converted to grayscale, and recalled to 512x512 pixels. We then generated 10 different contextual variations for each image. The changes were made using the object segmentation for each image obtained from the COCO object annotation masks and (for some conditions) replacing the background with different backgrounds based on the contextual manipulation conditions. The main manipulations per context type are as follows: (1) Full context: No manipulation, serving as the reference image with the object in a congruent context; (2) Incongruent context: Context swapped with a different (wrong) context; (3) No context: Context removed by swapping with gray pixels; (4) No object: Object removed by swapping with gray pixels; (5) Blurred context: Gaussian blur with kernel size 2 applied on the context; (6) Blurred object: Gaussian blur with kernel size 2 applied on the object; (7) Blurred incongruent boundary: Gaussian blur with kernel size 2 applied on the object-incongruent context boundary; (8) Minimal context: All context apart from the smallest bounding box around the object is removed; (9) Jigsaw context: 25x25 pixel context patches randomly shuffled around the object; and (10) Textured context: Context swapped with texture generated with Portilla & Simoncelli method^42^ (5 iterations) on the baseline image. Each of these context conditions was applied to 60 images, with 6 images per object category, resulting in a total of 600 images in the imageset.

#### Low-Level Image Features

For every image, we extracted a range of basic image features, such as object size, location and category, spectral mean and standard deviation(std), and contrast mean and standard deviation (std). The standard contrast metric for gray-scale images was used, calculated by the highest and lowest pixel values. The contrast standard deviation was derived from the pixel-wise standard deviation of the grayscale image. From the COCO object annotations, we determined the object size, represented in degrees of visual angle, as the fraction of the full image size (considering the full image was presented at 8 degrees) covered by the smallest bounding square around the object. The x and y coordinates, relative to the image, captured the object’s central position. Using the Fast Fourier Transform (FFT), we transformed the image in the spectral domain, and noted its spectral mean and standard deviation.

### Subjects

#### Human Participants

A total of 309 human subjects participated in the binary object discrimination tasks. Observers completed 5–10-min tasks through Amazon Mechanical Turk (MTurk), an online platform in which subjects could complete experiments for a payment of $15 CAD/hour. We confirm that this experimental protocol involving human participants was approved by and in concordance with the guidelines of the York University Human Participants Review Subcommittee.

#### Non human primates

The nonhuman subjects in our experiments were four adult male rhesus monkeys (*Macaca mulatta*). 2 of these monkeys (monkey M and monkey B), were trained with objects in congruent context and could perform the object discrimination tasks. The other 2 (monkey P, and monkey K) were naive to the discrmination task, and were only trained to passively fixate on the screen. All data were collected, and animal procedures were performed, in accordance with the NIH guidelines, the Massachusetts Institute of Technology Committee on Animal Care, and the guidelines of the Canadian Council on Animal Care on the use of laboratory animals and were also approved by the York University Animal Care Committee.

### Behavioral testing

#### Primate behavioral testing

##### Humans active binary object discrimination task

We collected large-scale psychophysical data from 309 subjects using Amazon Mechanical Turk (MTurk), an online crowdsourcing platform. The reliability of MTurk for psychophysical experiments has been previously validated by comparing online and in-lab results. Each trial began with a brief presentation (100 ms) of a sample image, selected from a set of 600 images. After a 100 ms blank gray screen, subjects were shown a choice screen displaying the target and distractor objects, similar to the procedure used in^22,23^. Subjects indicated their choice by touching the screen or clicking the mouse on the target object. No information regarding the sex of the participants was collected.

##### Macaques active binary object discrimination task

We measured monkey behavior from 2 male rhesus macaques. Images were presented on a 24-inch LCD monitor (1920 × 1080 at 60 Hz) positioned 42.5 cm in front of the animal. Monkeys were head fixed. Monkeys fixated a white cross (0.2^°^) for 300 ms to initiate a trial. The trial started with the presentation of a sample image (from a set of 640 images) for 100 ms. This was followed by a blank gray screen for 100 ms, after which the choice screen was shown containing a standard image of the target object (the correct choice) and a standard image of the distractor object. The monkey was allowed to view freely the choice objects for up to 1500 ms and indicated its final choice by holding fixation over the selected object for 400 ms. Trials were aborted if gaze was not held within ±2° of the central fixation dot during any point until the choice screen was shown. Prior to testing in the laboratory, monkeys were trained in their home-cages to perform the delayed match to sample tasks on the same object categories (but with a different set of images).

##### ANN behavioral testing

We evaluated eighteen ANN models, on the exact images shown to the macaques and humans. We focused on publicly available pre-trained PyTorch model architectures that have demonstrated significant success in computer vision benchmarks. Table 1 lists the models used and their characteristics.

To make these pre-trained models compatible with our specific 10-way object recognition task, we used the extracted features from each model for every stimulus, from the most IT-like layers (chosen based on BrainScore if that data was available, otherwise the most reasonable penultimate layer) shown in Table 1. To ensure consistency in results across the models, given the varying layer sizes for each, we standardized the dimension for every model down to 3,000 features. This was done by using Gaussian random projection with 3,000 components to project the full extracted features space on a randomly generated linear subspace in such a way that distances between the points are nearly preserved. We trained a multiclass SVM classifier using these scaled features (standard scaling) to calculate the cross validated probabilities for each object class (using 10 one-vs-all classifiers, 5 folds, 10 repetitions), mimicking the subjects’ active binary object discrimination task. All behavioral predictions from the decoder were for images where the object was not seen in any phase of the model training regardless of the surrounding context.

### Electrophysiological recording and data preprocessing

#### Passive Fixation Task

During the passive viewing task, monkeys fixated a white cross (0.2°) for 300 ms to initiate a trial. We then presented a sequence of 5 to 10 images, each ON for 100 ms followed by a 100 ms gray (background, ‘OFF’) blank screen. This was followed by fluid (water) reward and an inter-trial interval of 500 ms, followed by the next sequence. The animals (n = 2, male rhesus macaques) used in the passive fixation experiments study can be classified as “categorization task naive”, since they have not been explicitly trained to perform any object categorization tasks.

#### Eye Tracking

We monitored eye movements using video eye tracking (SR Research EyeLink 1000). Using operant conditioning and water reward, our 2 subjects were trained to fixate a central white square (0.2°) within a square fixation window that ranged from ±2°. At the start of each behavioral session, monkeys performed an eye-tracking calibration task by making a saccade to a range of spatial targets and maintaining fixation for 500 ms. Calibration was repeated if drift was noticed over the course of the session.

Real-time eye-tracking was employed to ensure that eye jitter did not exceed ±2°, otherwise the trial was aborted, and data discarded. Stimulus display and reward control were managed using the MWorks Software (https://mworks.github.io).

### Data Analyses

#### Behavioral Metrics

We developed two behavioral metrics, the hit rate at context level - B.C1 and more fine grained image level - B.I1 (as introduced in^22^). We obtained a biological or artificial signature for each system by applying each metric to its behavioral accuracies per image averaged across all trials. The one-versus-all context-level performance metric (B.C1) estimates the discriminability of all images of context category c, essentially pooling the accuracies across all images of context type c and all object/distractor pairs within. Because we tested 10 context categories, the resulting B.C1 signature has 10 independent values.

The one-versus-all image-level performance metric (B.I1) estimates the discriminability of each image, pooling across all distractors. Because we have an image test set of 600 images (60 per object, see above), the resulting B.I1 signature has 600 independent values. Given an image *i* of object category *o*, and all nine distractor objects (*d≠o*), we computed the average performance per image as: 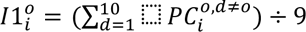, where *PC* (percent correct) is the fraction of correct responses for the binary task between object categories *o* and *d*. Considering every image *i_c_* of context type *c*, the B.C1 performance for each context type is the mean across the performance of all images (60 per context type): 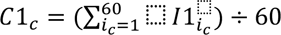

#### Human-monkey shared behavioral variance

To quantify the behavioral pattern similarity at a context and image level across humans and monkeys, we calculated the percent of shared behavioral variance (SV) for both signatures. The SV is obtained as the square of the correlation (Pearson’s R) of the pooled humans and pooled monkeys behavioral signature, corrected by the human and monkey signature internal consistency. This was repeated 20 times choosing 600 images with repetition (bootstrap). The ceiling estimates in Fig 7B and 7C show the full range for the 20 bootstrap values for C1 and I1 respectively.

#### Partial correlation analysis

To estimate the fraction of shared human-monkey variance that is explained by the models (including the Neural model), we calculated the partial correlation for the pooled humans and monkeys population - while controlling individually for each model. The partial correlation gives us the fraction of the primate shared variance that is independent of the model variance. The percentage of shared human-monkey variance explained by the model is then given by the formula:

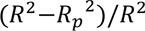, where 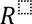 is the human-monkey Pearson correlation, *R_p_* is the human-monkey partial correlation, while controlling for the model (calculated as the product of the residuals of the model predictions).

The neural correction of the partial correlation is done by fitting a sigmoid extrapolation to an infinite number of neural trials (see **Fig 6D** inset for 122 neurons). Both 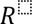 and *R_p_* are corrected by the (Spearman-Brown corrected) human and monkey split-half reliabilities 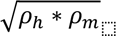, however, due to the normalization by *R*^2^, we did not need to account for the human or monkey noise.

#### Internal consistency

The reliability of each system (pooled human, monkey, and IT population) was assessed by calculating the trial split-half Spearman-Brown corrected correlation. For the pooled humans or monkeys, this was done by splitting all the accuracy trials per image in two halves, taking the mean for both halves (for each image), and computing the corrected Spearman correlation across all images for the two halves, repeated 100 times with different trial splits. The internal consistency for the decoding accuracy of the neural data was computed by calculating the decoding accuracy for each mean half of the neural trials and correlating the two obtained accuracies (across all images). The ceiling estimates shown in Fig 5B and 5D are the pooled human internal consistency, showing the full range of values (min-max).

### Statistical Analyses

For each statistical analysis, we first tested the normality of the data. We used the Lilliefors test assuming normal distribution, with a threshold 5% (normal distribution: p>0.05).

To test for statistical significance with a normal distribution of the data, a paired (monkey-human comparison) independent (comparing contextual variations) T-test was performed .This is a test for the null hypothesis that two samples have identical average (expected) values. The t(DOF)- statistic value quantifies the difference between the arithmetic means of the two samples. It is calculated as the mean of the difference of the two variables, divided by the standard error. The p-value quantifies the probability of observing as or more extreme values assuming the null hypothesis, that the samples are drawn from populations with the same population means, is true.

A Wilcoxon signed rank (paired variables) or ranksum (independent variables) was performed in case of a non-normal data distribution. The null hypothesis is that two (paired or independent respectively) samples come from the same distribution. In particular, it tests whether the distribution of the differences x -y is symmetric about zero. It is a non-parametric version of the T-test.

We chose the threshold 5% (p<0.05) to reject the null hypothesis for all tests.

## Acknowledgments

SD was supported by the Bertarelli Foundation Fellowship grant. HR is funded by CIHR Postdoctoral Fellowship. GK is supported by NIH R01EY026025. KK has been supported by funds from the Canada Foundation for Innovation (CFI), the Canada Research Chair Program, the Simons Foundation Autism Research Initiative (SFARI, 967073), the Canada First Research Excellence Funds (VISTA Program), and a Google Research Award.

## Supplementary Figures

**Fig S1.**
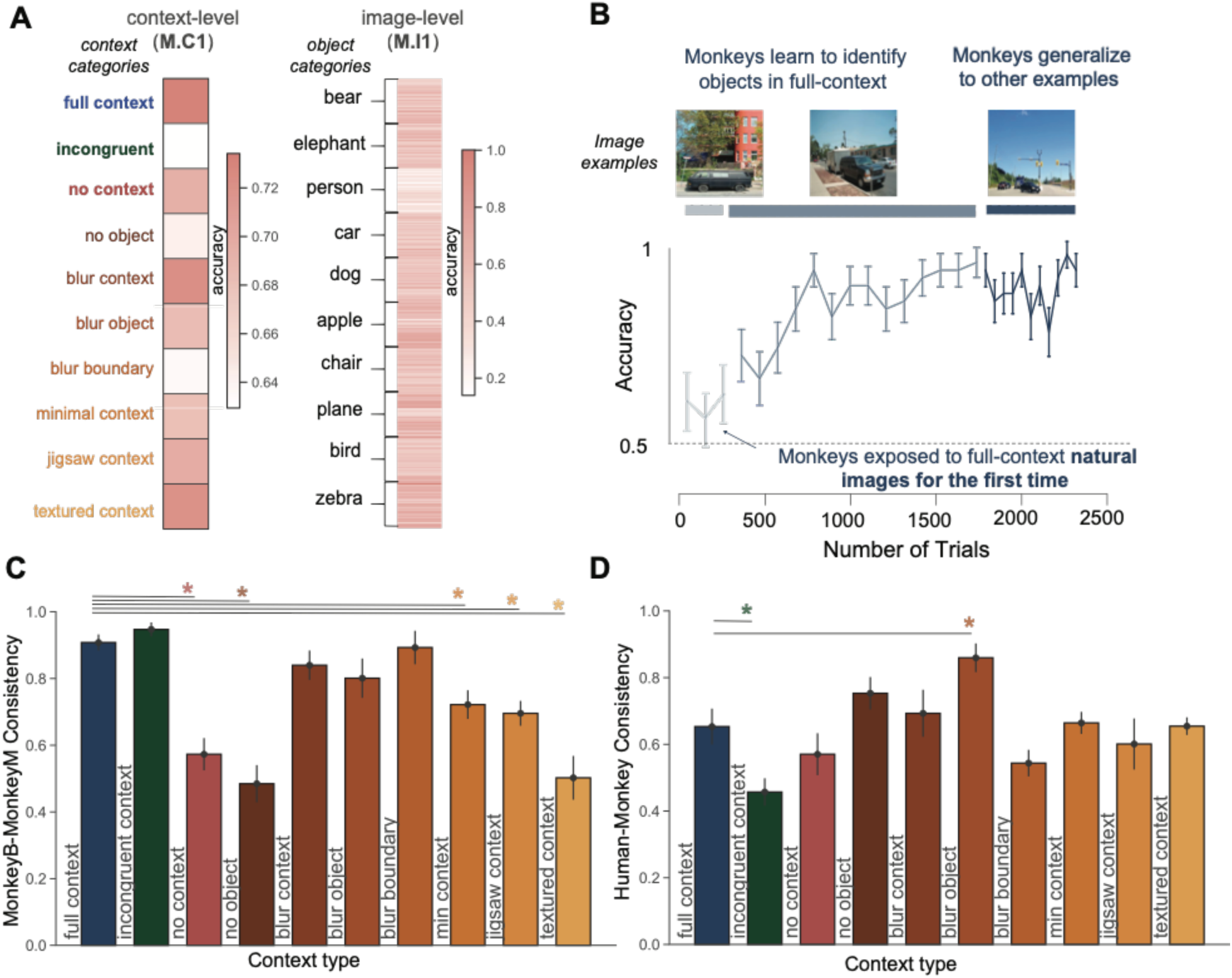
Monkeys’ behavioral signatures, context learning curve and image-level consistency per context type. **A.** Context-level (B.C1) and image-level (B.I1) behavioral signatures defined from the monkeys’ accuracy. Same as **Fig 2A**, but for the pooled monkeys. **B.** Training process for one macaque in the task shown in **Fig 1B** (chance = 0.5). The monkey, who was previously not exposed to images with objects in context, has low starting performance for full-context images (light blue curve). However, the monkey quickly learns to recognize images in full context (blue curve). Furthermore, this ability generalizes to new images (dark blue curve). Error-bars show the standard deviation across object categories. **C.** Each bar shows the corrected Pearson correlation between the two monkeys (monkey M and monkey B) for the images of a context category. Error-bars are standard errors across ten subsamples of images within a context category. Statistics are shown for full context compared to each other context variation (* denotes t-test, p<0.05). **D.** Similar to C but shows the corrected Pearson correlation between the pooled two monkeys and the pooled human population.

**Fig S2.**
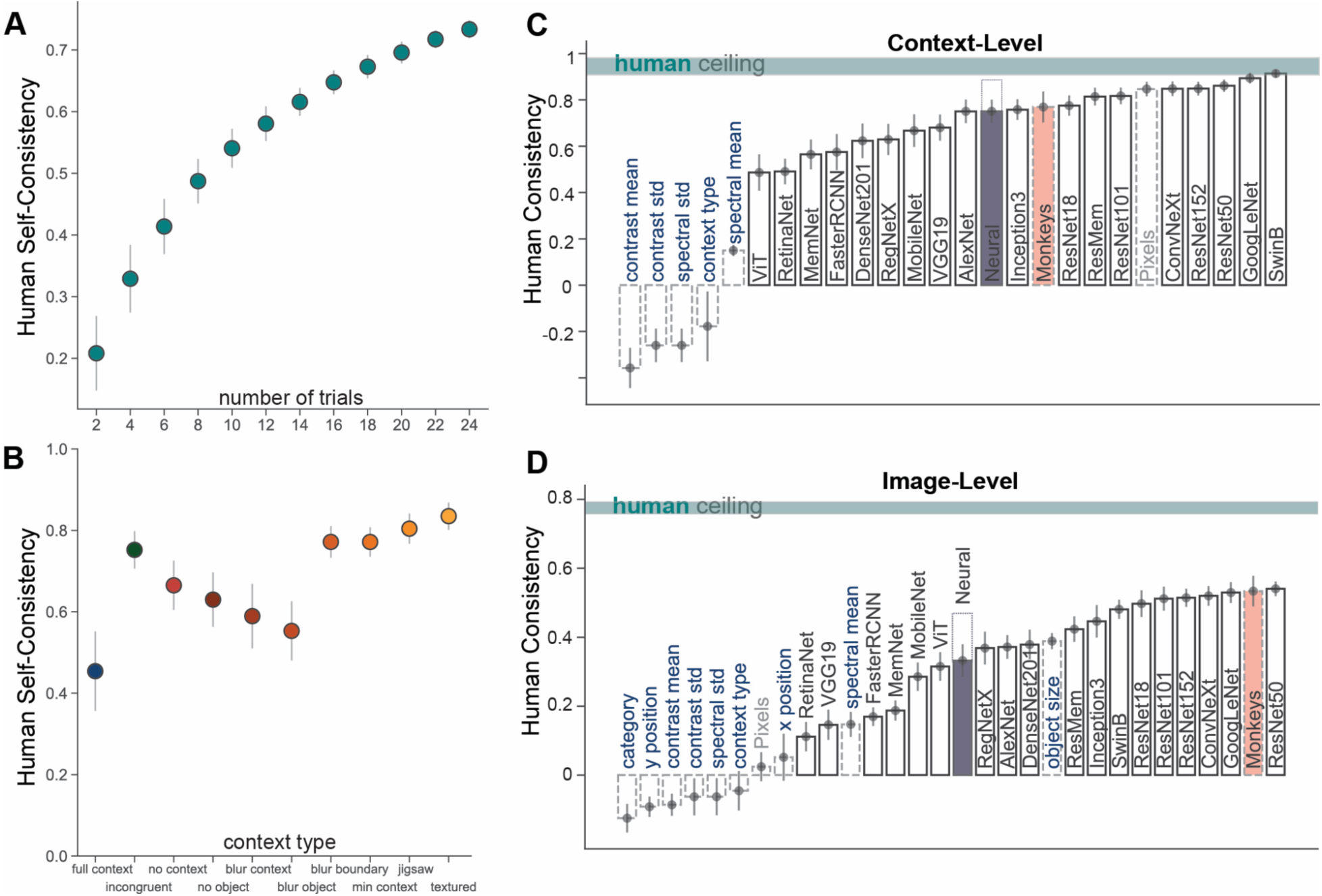
ANNs and IT population are consistent with human behavior at a context-level and only partially at an image-level. **A.** Human population self-consistency: Spearman-Brown corrected split-half correlation for increasing number of trials. The mean correlation for 100 different splits for each subset of trials, with standard deviation across the splits. **B.** Human population self-consistency (Spearman-Brown corrected split-half correlation) using all 24 trials per image, for images grouped by context category. The mean human internal reliability across 100 splits with standard deviation for each context category (color coding same as **Fig 1A**, and labeled on the x axis). **C.** The human consistency - Pearson R with the low-level features, ANN models and the Neural model at a context level. The low-level features are shown with light blue text (dashed bars). The mean human internal behavioral consistency ceiling is shown in gray, with standard deviation across different image subsamples. We show the noise corrected (by the split-half decoding consistency) neural consistency (purple bar), when using all the recorded reliable neural responses (122), the extrapolated consistency (to 4537 neurons, as in **Fig 6E**) is shown with a dashed bar on top. The noise corrected (by the monkey internal reliability) consistency with monkeys is shown in coral. **D.** Similar to C but showing the consistency at an image-level.

**Fig S3.**
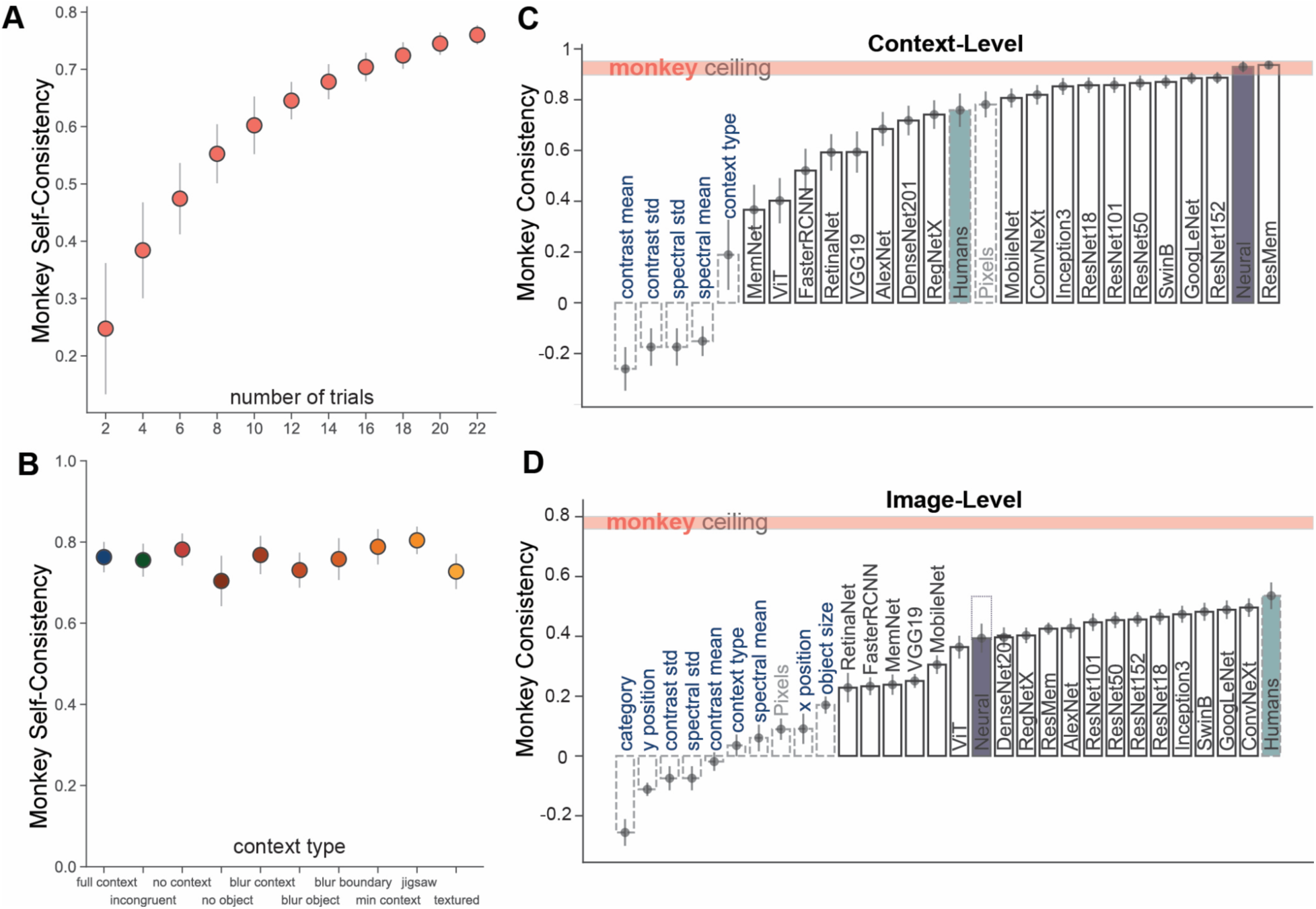
ANNs and IT population are consistent with monkey behavior at a context-level and only partially at an image-level. **A.** Monkey self-consistency: Spearman-Brown corrected split-half correlation for increasing number of trials. The mean correlation for 100 different splits for each subset of trials, with standard deviation across the splits. **B.** Monkey self-consistency (Spearman-Brown corrected split-half correlation) using all 22 trials per image, for images grouped by context category. The mean monkey internal reliability across 100 splits with standard deviation for each context category (color coding same as **Fig 1A**, and labeled on the x axis). **C.** The monkey consistency - Pearson R with the low-level features, ANN models and the Neural model at a context level. The low-level features are shown with light blue text (dashed bars). The mean monkey internal behavioral consistency ceiling is shown in gray, with standard deviation across different image subsamples. We are noting the noise corrected (by the split-half decoding consistency) neural consistency (purple) when using all the recorded reliable neural responses (122), the extrapolated consistency (to 4537 neurons, as in **Fig 6E**) is shown with a dashed bar on top (no extrapolation needed for context-level as the consistency is already within the monkey ceiling). The noise corrected (by the human internal reliability) consistency with humans is shown in teal. **D.** Similar to C but showing the consistency at an image-level.

**Fig S4.**
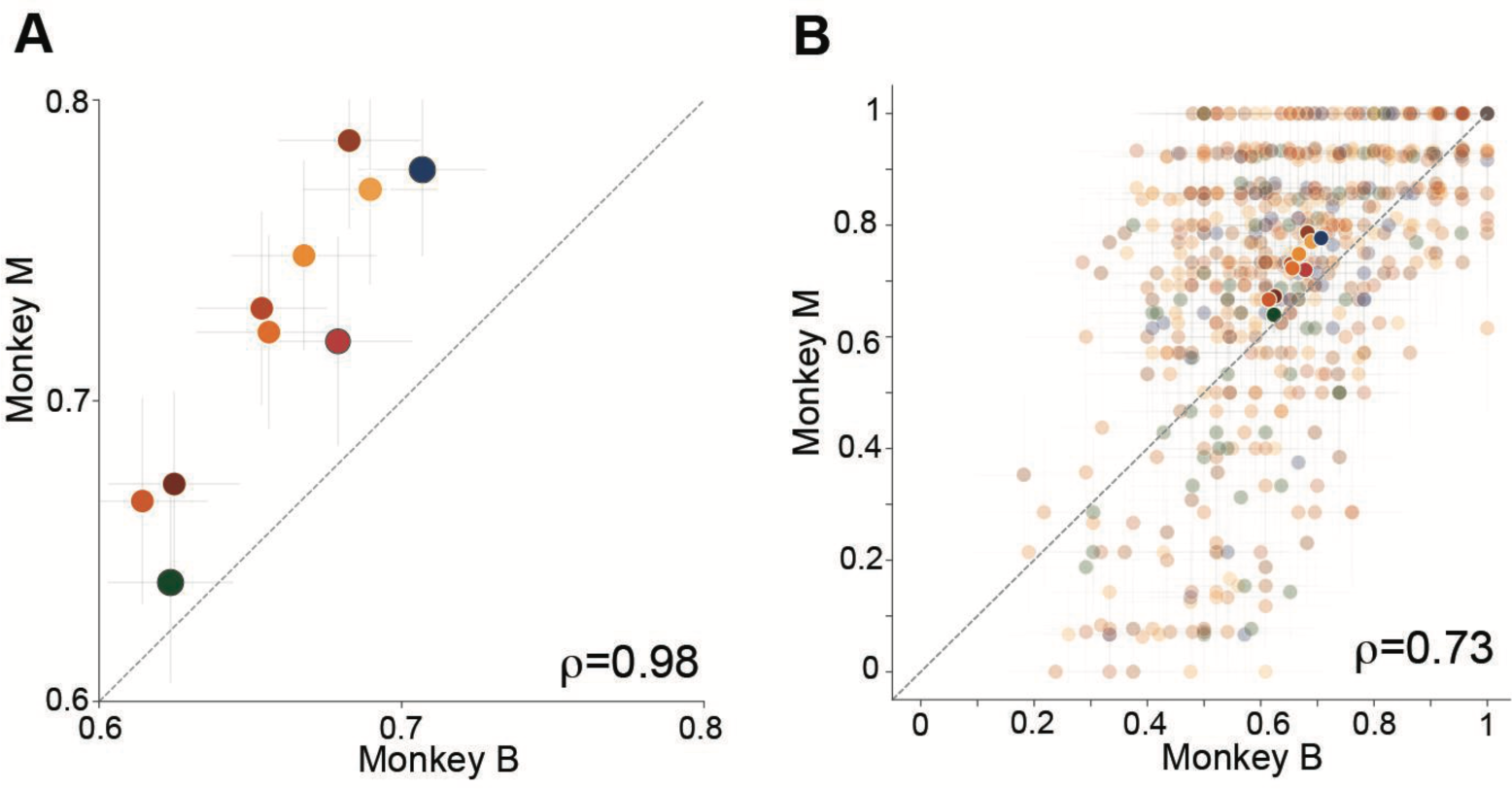
Two monkeys show similar (but not identical) context-driven behavioral changes. **A**. Context-level (B.C1) correlation between the two monkeys. Each point represents the mean accuracy for a contextual variation with standard error across images of that context type (colors as in **Fig 1A**, Monkey B mean 0.66±0.03, Monkey M mean 0.72±0.05). The three main context types: full (blue), incongruent (green) and no context (red), are shown with a black stride. The value ρ indicates the noise corrected correlation coefficient (Pearson R). **B.** Image-level correlation (B.I1) for the two macaques, each low opacity point shows the performance(accuracy) for an image with standard error across trials, the higher opacity points are the B.C1 mean (from A), colors map to context types as defined in **Fig 1A**.

**Fig S5.**
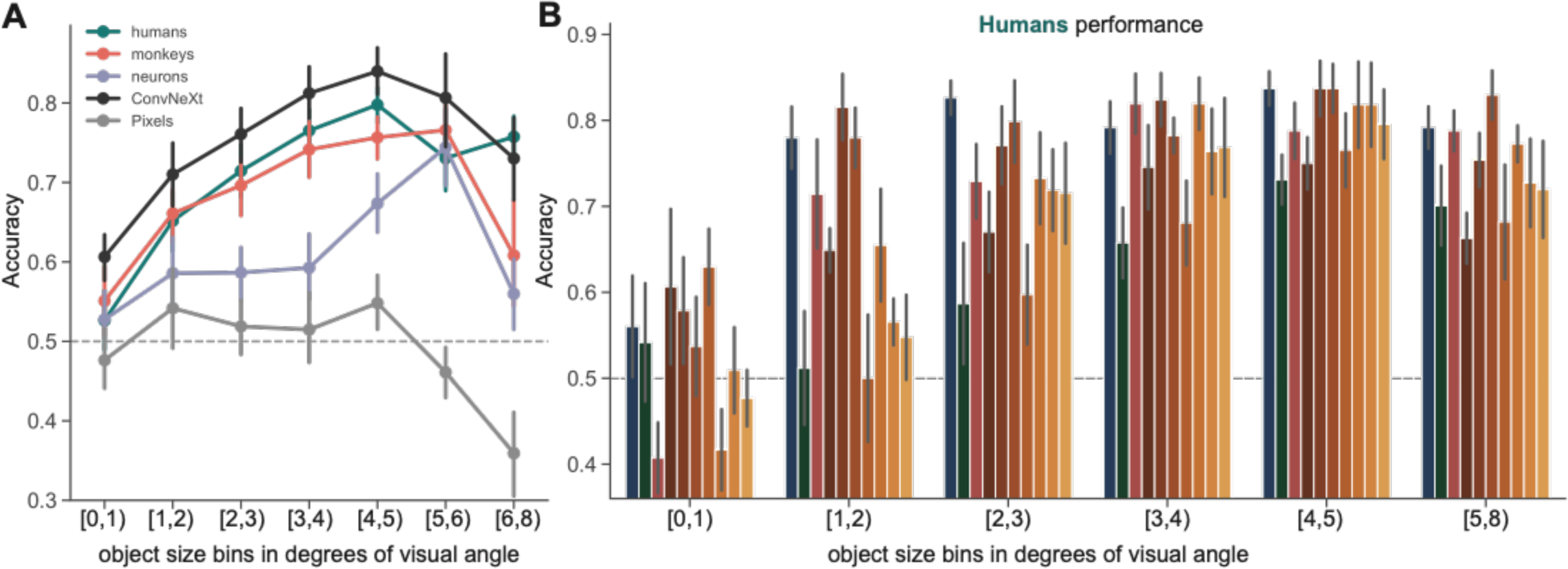
Object size predicts primate and ANN average accuracy. **A.** Average image-level accuracy (for all images) grouped in bins based on the object size (in degrees of visual angle, the full image is 8 degrees), with standard error across images in each bin. The performance is shown for humans (teal), pooled monkeys (coral), IT population (purple), Pixels (gray) and ConvNeXt (black) - the best model explaining the highest fraction of the human-monkey shared image-level behavioral variance (see Fig 7C). Chance level accuracy (0.5) is noted with the dashed gray line. The size bins are labeled with the minimum size and max size (not included) of the bin (eg. the size bin [1,2) contains all images where the object size is greater or equal to 1 degree and smaller than 2 degrees). **B.** Human accuracy for each object size grouped by context category (color coding for context type from Fig 1), with standard error across images in each bin. The human data in part A (teal) is the average over all context conditions shown in part B.

**Fig S6.**
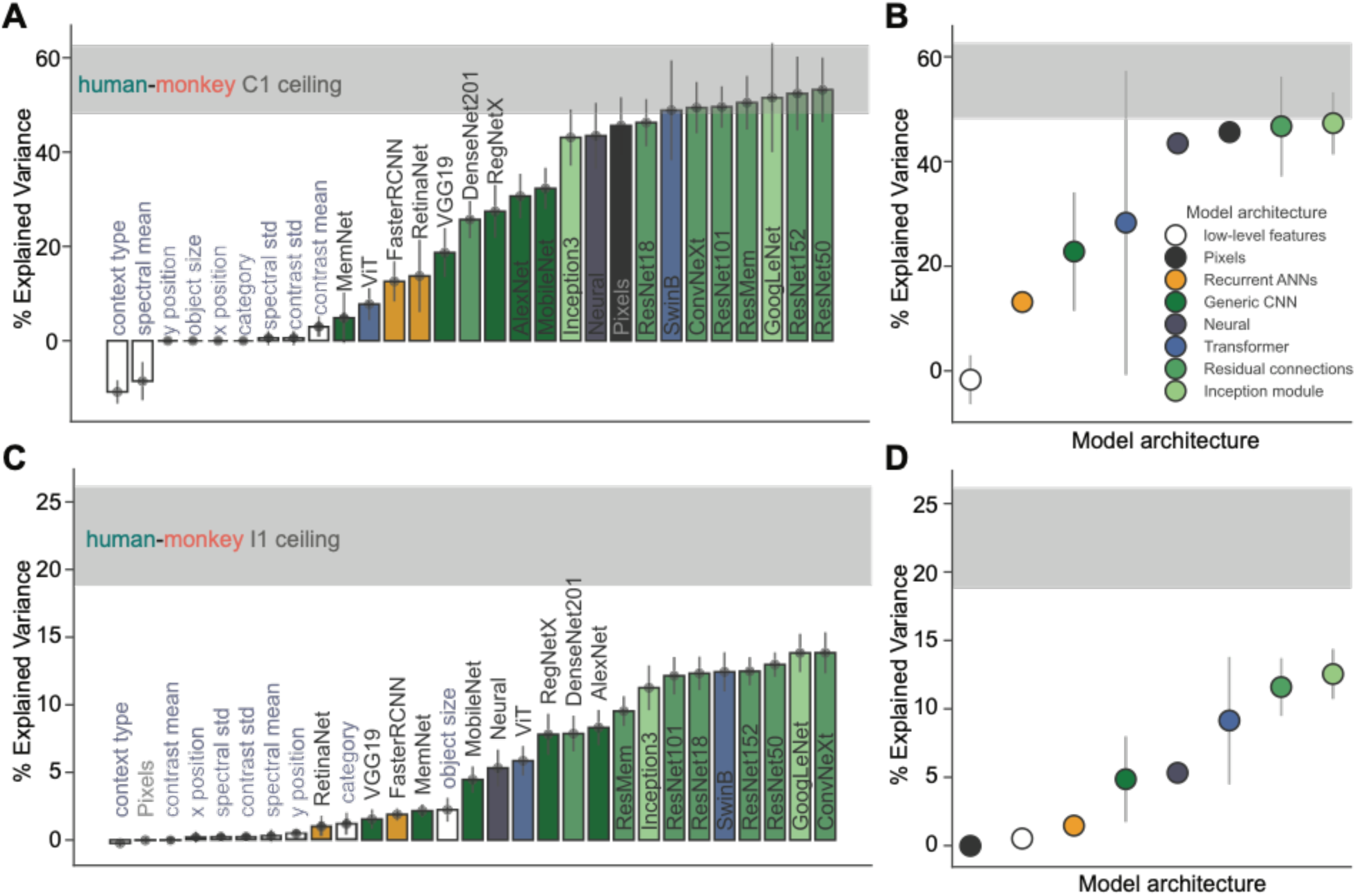
ANN model architecture effects on explained human-monkey shared variance. **A.** Shows the same models as **Fig 7**, but the bars are color coded for the model architecture (see legend in B). Green is used for CNNs (with subgroups: models with Inception modules and residual connections), blue for visual transformers and yellow for recurrent neural networks. **B.** The model performance, grouped by model architecture, with the standard deviation across models. CNN models with residual or inception blocks share the most of the shared human-monkey variance. **C.** Same as A but for image-level EV. **D.** Same as B but for image-level EV.

**Fig S7.**
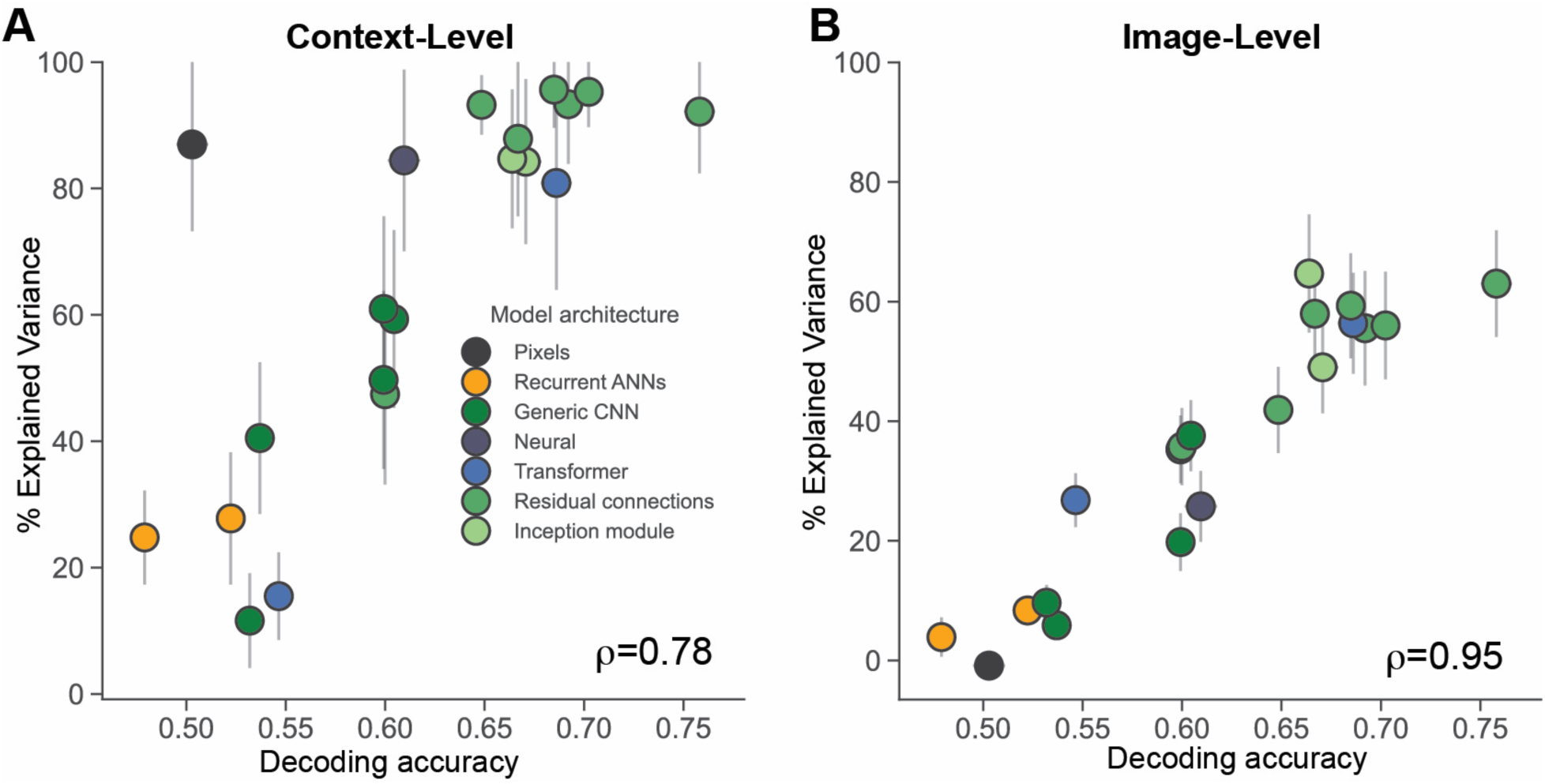
The models’ decoding accuracy predicts the fraction of explained shared human-monkey variance. **A.** The percent of variance explained by each model from the shared human-monkey variance at a context-level as a function of the mean image decoding accuracy for each model. Each point is a different ANN model (Neural model in purple, Pixels in black), color coded by model architecture (as in **Fig S6**, see legend and Table 1). The y axis shows the normalized % EV (by the human-monkey shared variance ceiling) with standard deviation across image subsamples for each model. The x axis shows the mean decoding accuracy across all images (with standard error, x error bars are smaller than the points). ⍴ notes the Pearson correlation between the accuracy and EV across models. **B.** Same as A but showing the %EV as a function of the decoding accuracy at an image-level.

**Fig S8.**
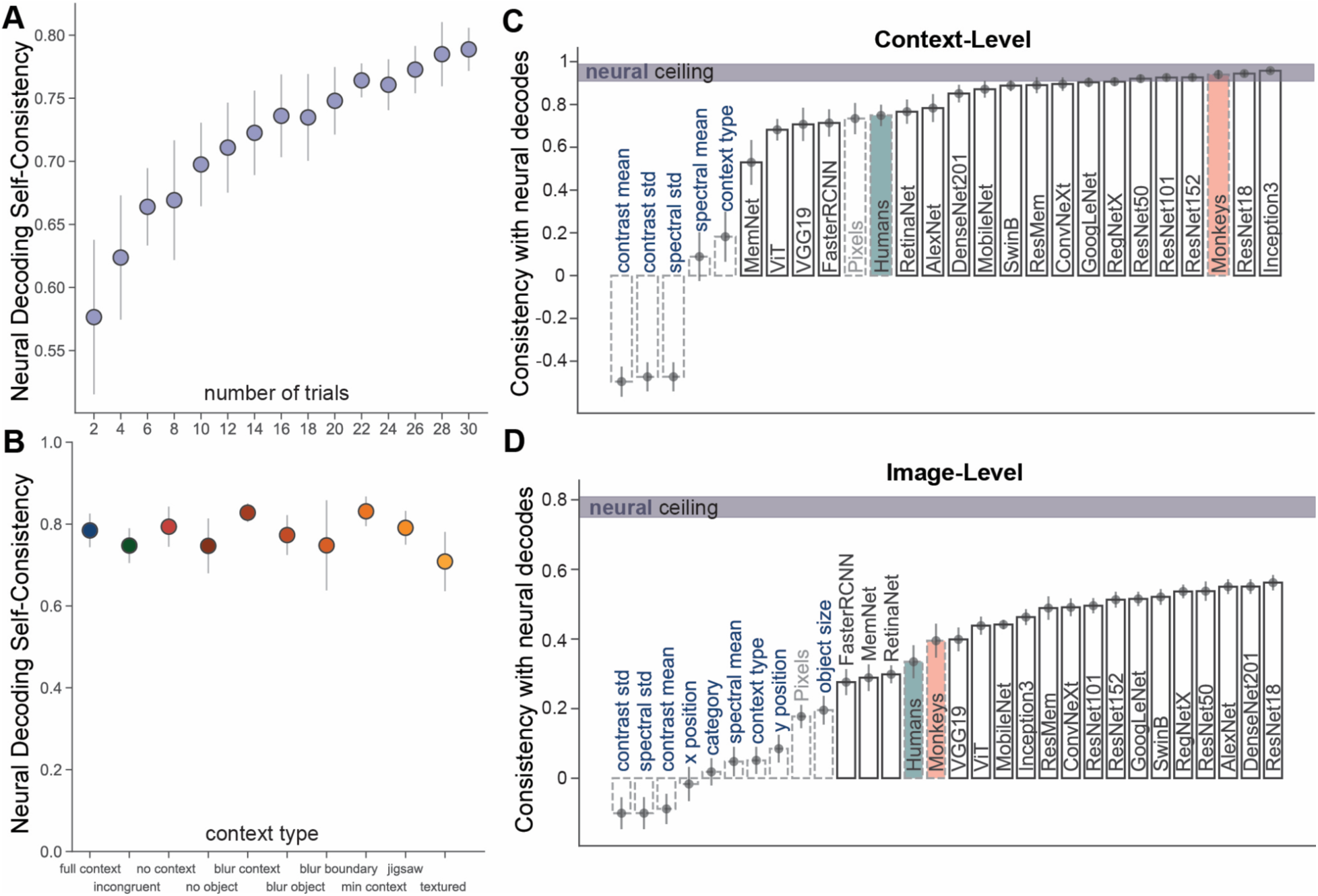
ANNs and primates are consistent with context-naive IT decoded behavior at a context-level and only partially at an image-level. **A.** Neural decoding accuracy self-consistency: Spearman-Brown corrected split-half correlation of the decoding accuracy across all images for increasing number of neural trials used for decoding (n=122 neural sites used). The mean correlation for 20 different splits for each subset of trials, with standard deviation across the splits. **B.** Neural decoding accuracy self-consistency (Spearman-Brown corrected split-half correlation) using all 30 neural trials per image, per neuron (for the 122 neurons), for images grouped by context category. The mean neural decoding internal reliability across 20 splits with standard deviation for each context category (color coding same as **Fig 1A**, and labeled on the x axis). **C.** The consistency with neural decode based predictions (Pearson R) with the low-level features, ANN models, pooled humans and monkeys behavioral accuracy at a context-level. The low-level features are shown with light blue text (dashed bars). The mean neural internal behavioral consistency ceiling is shown in gray (split-half decoding reliability), with standard deviation across different image subsamples. We are noting the (internal reliability corrected) human (teal) and monkey (coral) consistency. **D.** Similar to C but showing the consistency at an image-level.

**Fig S9.**
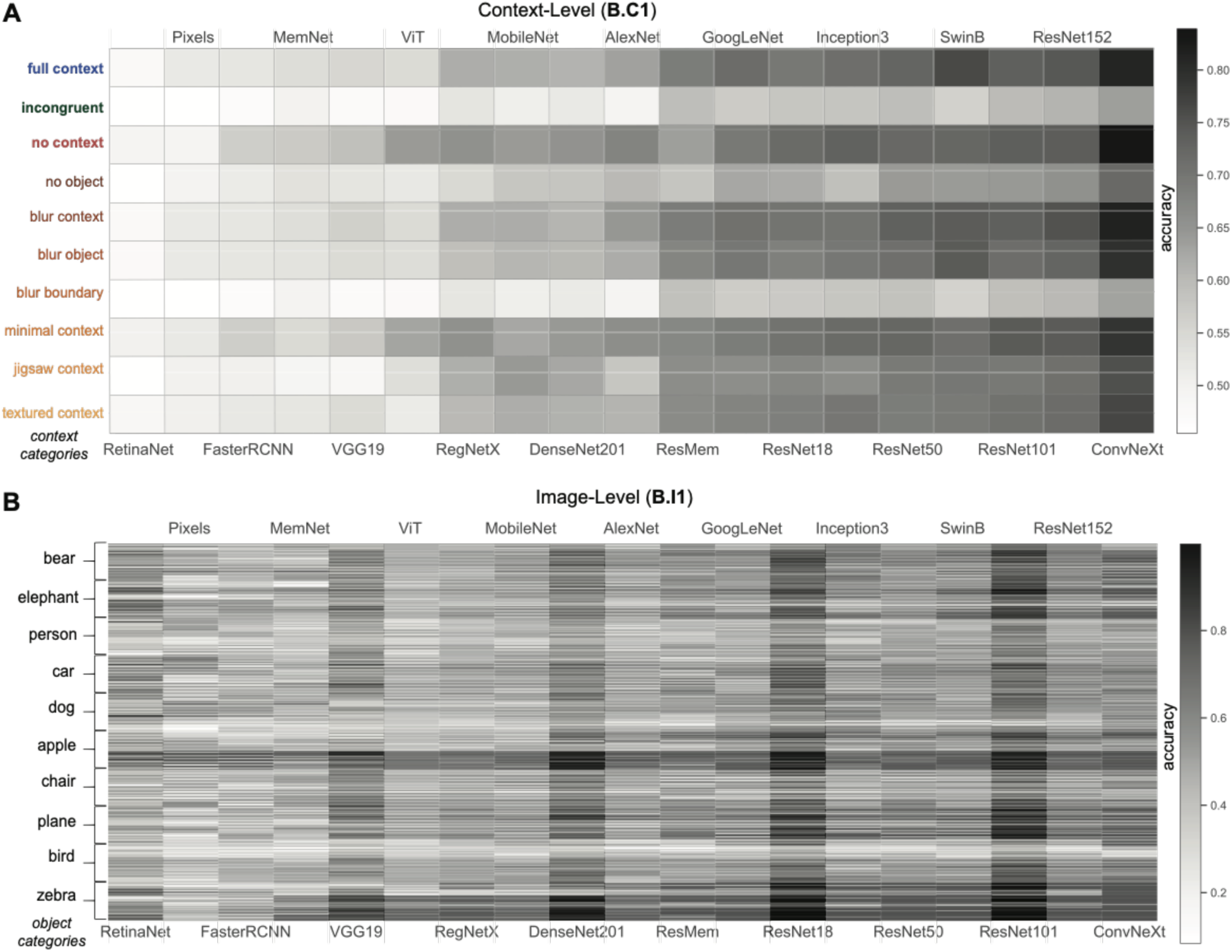
Pixels and ANN models behavioral signatures. **A.** Context-level behavioral signature (B.C1) for each model (column), sorted by their overall average decoding accuracy - least accurate (left) to most accurate (right). The average accuracy is shown for each context type (row) with the color indicating the accuracy (increasing from white to black, see colorbar on the right). **B.** Similar to A but showing the image-level signature for each model (B.I1), each line represents the image accuracy averaged across trials and distractors, with the rows sorted and grouped by object category (see on the left).

